# Contributions of distinct attention mechanisms to saccadic choices in a gamified, dynamic environment

**DOI:** 10.1101/2025.01.25.634882

**Authors:** Evan A Kattner, Terrence R Stanford, Emilio Salinas

**Affiliations:** Department of Translational Neuroscience, Wake Forest University School of Medicine, 1 Medical Center Blvd., Winston-Salem, NC 27157-1010, USA

## Abstract

Visuospatial attention is key for parsing visual information and selecting targets to look at. Based on regimented laboratory tasks, it is now well established that three types of mechanism determine when and where attention is deployed; these are stimulus-driven (exogenous), goal-driven (endogenous), and history-driven (reflecting recent experience). It is unclear, however, how these distinct attentional signals interact and contribute in visual environments that are more akin to natural scanning, when stimuli may change rapidly and no fixation requirements are imposed. Here, we investigate this via a gamified task in which participants (male and female) make continuous saccadic choices at a rapid pace — and yet, perceptual performance can be accurately tracked over time as the choice process unfolds. The results reveal unequivocal markers of exogenous capture toward salient stimuli; endogenous guidance toward valuable targets and relevant locations; and history-driven effects, which produce large, involuntary modulations in processing capacity. Under dynamic conditions, success probability is dictated by temporally precise interplay between different forms of spatial attention, with recent history making a particularly prominent contribution.

**Significance Statement:** Visuospatial attention comprises a collection of mental mechanisms that allow us to focus on (or look at) specific objects or parts of space and ignore others. The next target to be inspected is generally selected based on how much it stands out (salience), its relevance to current goals, and recent experience. We designed a gamified visual scanning task in which all such forms of attentional control interact rapidly, more akin to real life situations (e.g., driving through traffic). Each mechanism affected in characteristic ways the probability that participants would look to the correct target at each moment in time. Most notably, we found that the history of recently seen stimuli determines visual processing capacity much more strongly than previously thought.

## Introduction

Day-to-day we experience a complex visual world, and navigating it successfully requires a system by which to prioritize information and select relevant parts. Visuospatial attention is the cognitive capacity to do precisely that, i.e., direct processing resources to areas of visual space that are important while ignoring areas that are not.

Much is known about the specialized mechanisms that enable the deployment of attention, either overtly (via eye movements) or covertly (with gaze fixed), to specific points in space (Itti and Koch, 2001; Corbetta and Shulman, 2002; Theeuwes, 2010; Carrasco, 2011; Wolfe and Horowitz, 2017). Bottom-up or exogenous mechanisms draw attention rapidly, transiently, and involuntarily toward physically salient stimuli. In contrast, top-down or endogenous mechanisms are slower and require effort, but permit the voluntary and sustained deployment of attention to potentially goal-relevant objects or locations regardless of physical salience. In addition to this traditional dichotomy, it is now widely recognized that recent experience also plays a critical role in determining the effective strength of visual features or locations (Fecteau and Munoz, 2003; Awh et al., 2012; Theeuwes, 2019; Anderson et al., 2021). Lingering biases due to prior history take many forms, depending on past stimuli, past actions, past rewards, etc (Maljkovic and Nakayama, 1994; Failing and Theeuwes, 2018; Jiang and Sisk, 2019; Kim and Anderson, 2019; Anderson et al., 2021). In primates, whenever oculomotor circuits redirect the eyes to a new fixation target (every 200–250 ms), the underlying selection process is heavily steered by the three types of mechanism just listed: stimulus-driven, goal-driven, and history-driven (Maunsell, 2015; Moore and Zirnsak, 2017; Westerberg and Schall, 2021).

Although these mechanisms have been studied extensively, prior work has largely relied on segmented laboratory tasks with prolonged fixation requirements. Thus, it is unclear what their contributions would be like under more dynamic and less constrained viewing conditions; say, while playing basketball or driving through traffic. Here, we investigate visuospatial attention in a laboratory experiment that takes a few steps toward such richness. Why is this important? More ecologically realistic situations may reveal not only subtle aspects of behavior that might easily go unnoticed, but also large, unexpected effects on visual processing (Camerer and Mobbs, 2017; Snow and Culham, 2021).

With less control, quantifying performance becomes more challenging. To strike a balance, we designed a new paradigm that combines two elements, richer dynamics and so called “urgent” choice tasks (Stanford and Salinas, 2021). Urgent tasks impose time presssure to yield a more variable and realistic interaction between motor plans and perceptual information than traditional, non-urgent tasks. The upshot is that it becomes possible to construct a unique psychophysical curve describing how a choice is elaborated as a function of time — and such “tachometric” curve reveals clear signatures of exogenous, endogenous, and history-based contributions to performance (Salinas et al., 2019; Goldstein et al., 2022, 2024; Oor et al., 2023, 2025)

Thus, we developed SpotChase, a gamified task that, rather than discrete trials, presents a continuous stream of choices akin to pro- and antisaccades. Instead of fixating and responding based on a rigidly timed script, participants make a series of perceptual judgements and eye movements at a (fast) pace that they control, aiming to obtain a high score across numerous choices in accordance with simple rules for accruing points. This produces scanning behavior that is more rapid, interactive, and variable than in standard segmented tasks. Critically, though, SpotChase retains sufficient stimulus control to quantify visuomotor performance rigorously via tachometric curves.

Using SpotChase, we found unequivocal, temporally segregated behavioral signatures of exogenous and endogenous attention similar to those reported previously (Salinas et al., 2019; Goldstein et al., 2022, 2024; Oor et al., 2023). But in this case, selection history effects were most remarkable: they resulted in powerful, involuntary modulation of both the early (exogenously dominated) and later (endogenously dominated) phases of the target selection process, and accounted for wide differences in perceptual capacity between individual participants.

## Results

To study the contributions of different attention mechanisms to visuomotor performance under dynamic conditions, we ran several variants of SpotChase. We first describe the basic paradigm (base version), corresponding psychophysical results, and their relation to oculomotor parameters. Then, in subsequent subsections, we describe results from four task variants designed to characterize the impact on performance of cue salience (akin to exogenous attention), prior knowledge of the cue location (akin to covert attention), and task statistics and selection history.

### SpotChase: a dynamic choice task

Participants completed “runs” of SpotChase, 90 second periods of continuous performance of choices akin to pro- and antisaccades, but guided by an overall goal of accruing as many points as possible by the end of each run. Short videos that recreate real-time performance in SpotChase based on recorded data from two example participants are available at https://doi.org/10.5281/zenodo.17137751.

Every run began with an initial fixation on a red spot at the top-center of the screen (Fig. 1a, Fix). This was followed by presentation of two diametrically positioned yellow targets to the left and right of the center (Fig. 1a, Targets). After a variable interval (typically 50–300 ms), one of the two yellow targets switched to red or blue (Fig. 1a, Cue). After this, the display did not change until the participant made a choice by looking at one of the two colored spots (Fig. 1a, Saccade). Then, following a brief post-choice interval (150 ms), a new pair of yellow targets was presented, this time arranged vertically. This marked the start of a new choice cycle consisting of the same event sequence: Targets, Cue, and Saccade. Horizontal and vertical targets alternated throughout each run, allowing for seamless transitions between choices.

**Figure 1.**
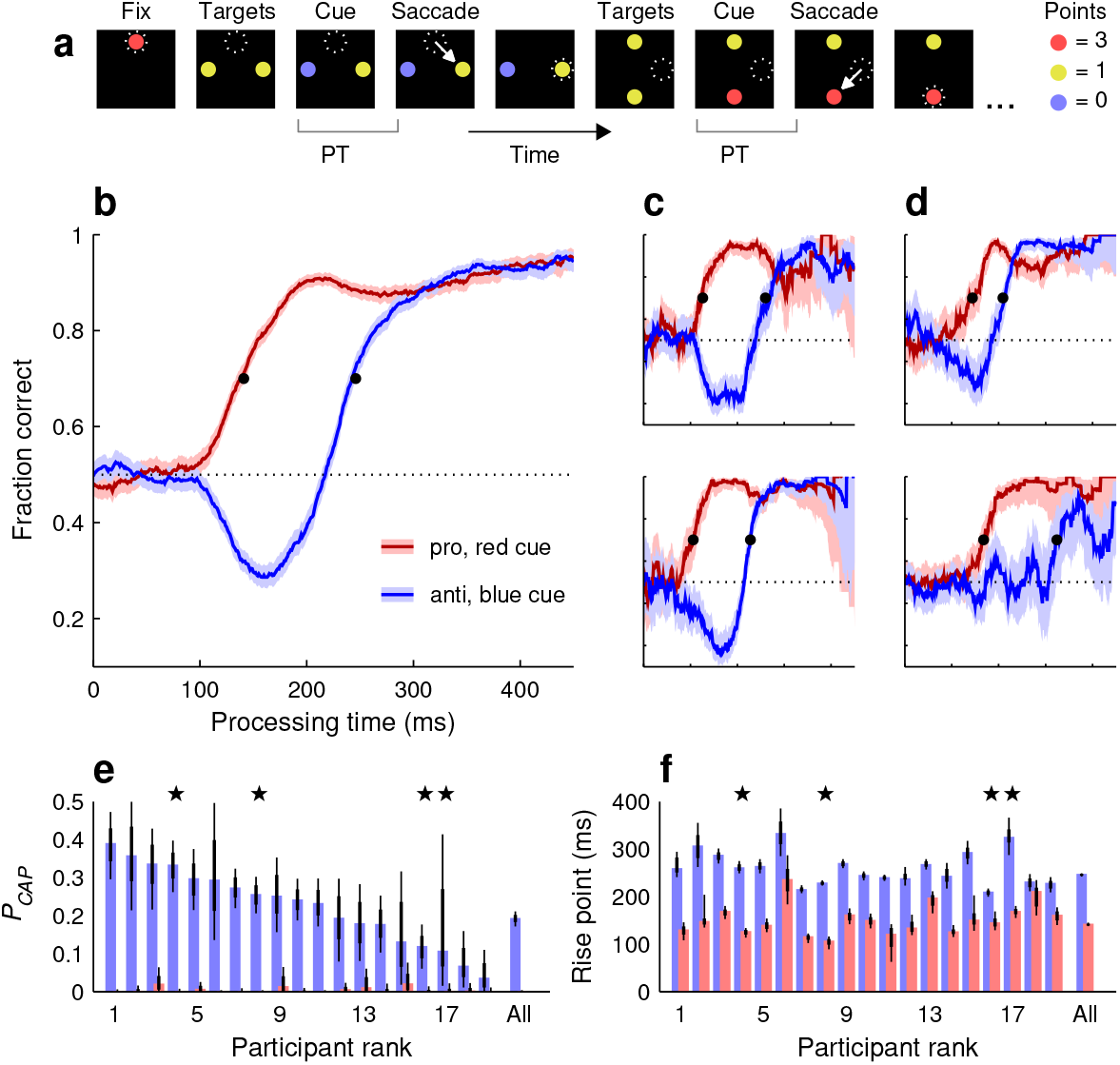
Performance in the base version of SpotChase. **a**, Sequence of events during the task. Each run begins with gaze (dotted circles) fixated on a red dot (Fix). This is followed by the appearance of two yellow targets aligned horizontally (Targets). After a variable time, one target switches to blue or red (Cue), prompting the participant to make a saccadic choice based on the point values shown. Once a choice is made (Saccade; arrows), vertical targets appear and the choice process starts anew. Thereafter, horizontal and vertical targets alternate until the run (90 s) ends. The processing time (PT) is the interval between cue onset and saccade onset. **b**, Tachometric curves for red-cue, or pro choices (red trace) and blue-cue, or anti choices (blue trace) combined across 19 participants. Shaded ribbons indicate 95% confidence intervals (CIs) for the mean value across choices. Horizontal dotted line marks chance performance. Black dots indicate rise points. **c, d**, Tachometric curves from individual participants with high (**c**) or low (**d**) *P*_*CAP*_ values. Same conventions as in **b**. Each curve from each participant is based on ∼1,100 choices. **e**, Probability of capture (*P*_*CAP*_) across participants. Bars indicate values for pro (red) and anti (blue) choices. Thin and thick lines indicate 95% and 68% CIs, respectively. Participants are sorted by their effect magnitude. Stars mark the example participants in **c** (pair on left) and **d** (pair on right). **f**, Curve rise points across participants. Same conventions and participant order as in **e**.

The participants aimed to accrue as many points as possible after each run, with points assigned according to the color of each chosen spot (Methods). Looking at a red, yellow, or blue target yielded 3, 1, or 0 points, respectively. In the base version of the task, red and blue cues were randomly interleaved with equal probability at all locations and had the same luminance as the yellow targets. Nevertheless, both cue stimuli were salient because of their sudden color onset. This, together with the scoring convention, established choice types similar to those of traditional prosaccade and antisaccade tasks, in that, to maximize their score, participants would direct their gaze towards a salient high-value (red) target or away from a salient low-value (blue) target.

It is worth emphasizing a few key differences between the continuous dynamic of SpotChase and traditional, trial-based oculomotor tasks. (1) After fixation on the red spot at the beginning of each run, no further fixation requirements were enforced. Participants learned to make choices efficiently, directing saccades mostly to the choice targets (Methods). (2) Feedback about performance was useful, but during data collection (after a few preliminary practice runs; Methods), it was provided only at the end of each run in the form of a score. (3) The pace of the task was controlled by the participants, who were free to make choice saccades either before, shortly after, or well after each cue onset, increasing their probability of success accordingly. However, the longer they waited to respond, the fewer choices they could make. In order to maximize the score, it was best to go at a relatively fast pace at which guesses were frequent but more choices became available.

It is also important to note that SpotChase is similar to trial-based urgent tasks in one critical respect: for any given choice, it permits accurate measurement of the interval between cue onset and saccade onset, which is the maximum amount of time available for viewing and interpreting the cue. This quantity, which is called the processing time (**PT**; Becker and Jürgens, 1979), dictates performance in urgent tasks (Salinas et al., 2019; Poth, 2021; Stanford and Salinas, 2021; Goldstein et al., 2022, 2024; Oor et al., 2023, 2025; Krause and Poth, 2025), and we expected it would do so in SpotChase as well. The rationale is straightforward: the longer the PT, the higher the likelihood that the choice is guided by the cue information and, in turn, the greater the probability of choosing the higher-value target. As a result, the tachometric curve, the curve that describes choice accuracy as a function of PT, is the critical behavioral metric to consider (for further detail about its advantages, see Stanford et al., 2010; Shankar et al., 2011; Salinas et al., 2014; Stanford and Salinas, 2021).

### SpotChase reveals signatures of exogenous and endogenous mechanisms

Nineteen participants performed the base version of Spotchase (Methods), on average generating 2.1 choices per second (range: 1.6–2.8 choices/s). For reference, participants performed approximately 0.7 choices/s in prior studies with comparable trial-based tasks (Goldstein et al., 2022, 2024). Tachometric curves were used to quantify performance (Methods). That is, choices were sorted into PT bins and, for each bin, the fraction of correct outcomes was calculated — where ‘correct’ means that the participant looked at the higher-value target on offer.

Separate tachometric curves were generated for red- and blue-cue choices (Fig. 1b-d) which, as noted above, correspond to situations in which a prosaccade (toward a cue stimulus) or an antisaccade (away from a cue stimulus) is requested. Indeed, the resulting curves revealed characteristic trends similar to those seen in urgent, trial-based versions of the pro- and antisaccade tasks (Salinas et al., 2019; Goldstein et al., 2022). During short-PT choices (PT ≲ 90 ms), participants had little or no time to view the cue information and accuracy remained at chance level (50% correct) regardless of cue color. Responses in this range were uninformed and correspond to guesses. At the opposite extreme (PT ≳ 300 ms), both curves achieved asymptotic accuracy levels near 90% correct, indicating that with sufficient cue viewing time participants consistently made informed choices in accordance with the goal of the task. Critically, in the transition between uninformed and informed choices, the salient cue onset gave rise to involuntary oculomotor capture, resulting in opposing deflections in performance for the two choice types. In anti choices (Fig. 1b-d, blue traces), for which salience-driven and goal-driven signals pointed in opposite directions, performance first dropped well below chance (due to exogenous capture) before rising steadily (due to endogenous guidance). In contrast, in pro choices (Fig. 1b-d, red traces), the exogenous and endogenous signals were aligned, leading to a rapid, monotonic upswing in performance. These trends were evident for the pooled data (Fig. 1b) as well as for individual participants (Fig. 1c, d).

Two measures were used to quantify these effects. To characterize the strength of exogenous capture in anti choices, we computed the probability of capture, or *P*_*CAP*_ (Fig. 1e), which is the proportion of incorrect responses beyond that expected by chance (Methods). In essence, *P*_*CAP*_ measures how much the tachometric curve dips below the chance line. This quantity was well above zero for the pooled data (*P*_*CAP*_ = 0.19 in [0.17, 0.21], 95% CI, *p <* 0.0001 from resampling test; Methods) and for 17 of the 19 individual participants (*P*_*CAP*_ range was 0.09–0.38 for *n* = 17 participants with *p <* 0.0001; *p >* 0.9 for *n* = 2 participants; Fig. 1e, blue bars). The exogenous capture toward the blue cue was highly robust, even though it acted briefly (for PTs in the 100– 225 ms range, approximately) and even though the yellow, red, and blue spots were isoluminant (Methods).

By way of control, we also computed *P*_*CAP*_ for the pro tachometric curves, which in general rise steadily above 50% correct. In this case, all the values were close to zero and none were larger than expected by chance (Fig. 1e, red bars). Thus, as anticipated, no capture was detected during pro choices.

In addition, to measure when the increase toward asymptotic accuracy occurred, we calculated the rise point of each tachometric curve (Fig. 1f), which was defined as the PT at which the fraction correct first exceeded 0.7 (Methods). By this measure, performance in pro choices rose much earlier (141 ms in [138, 145], 95% CI) than in anti choices (246 ms in [242, 250], 95% CI) based on the pooled data. The difference of 105 ms was consistent across individual participants, in that it was always positive (range: 20–197 ms, *n* = 19) and statistically large (no overlap between 95% CIs for pro and anti rise points) in all but one of them (Fig. 1f). Thus, to produce a correct informed saccade, processing the anti (blue) cue typically required about 100 ms more than processing the pro (red) cue.

### Relationship between perceptual and oculomotor performance

It is important to emphasize that the tachometric curves just discussed (Fig. 1b–d) are a quantitative measure of *perceptual* performance (we use this term as a shorthand that includes perceptual and closely allied cognitive processes, such as those necessary for mapping a cue color onto an appropriate target location). This is because the curves are strongly decoupled from motor readiness. To appreciate this, consider how urgency would impact the mean fraction correct over a full run. If the participants, say, increased their pace (rate of choices per second), one would expect a drop in fraction correct in accordance with the speed-accuracy tradeoff (Wickelgren, 1977; Chittka et al., 2009). By contrast, the tachometric curve is largely impervious to such tradeoff (Stanford et al., 2010; Salinas et al., 2014, 2019). If urgency increased, the left part of the curve would be sampled more heavily, so more guesses would be made and overall accuracy would drop — but the shape of the curve would not change.

To verify such decoupling of perceptual and motor processes in SpotChase, pro and anti tachometric curves were compared across urgency conditions. For each choice, the reaction time (**RT**) was measured from the onset of the yellow targets to the onset of the choice saccade (Methods). The data from each participant were then split by the median RT and aggregated across partici-pants into two subsets, fast and slow. There were vast differences between the two resulting data subsets, not only in RT (mean of fast RTs: 211 ms, slow RTs: 428 ms) but also, as expected, in overall fraction correct (for fast RTs: 0.55; for slow: 0.74). In spite of this, as a function of PT, performance for each choice type was essentially the same in the two conditions (Fig. 2a).

**Figure 2.**
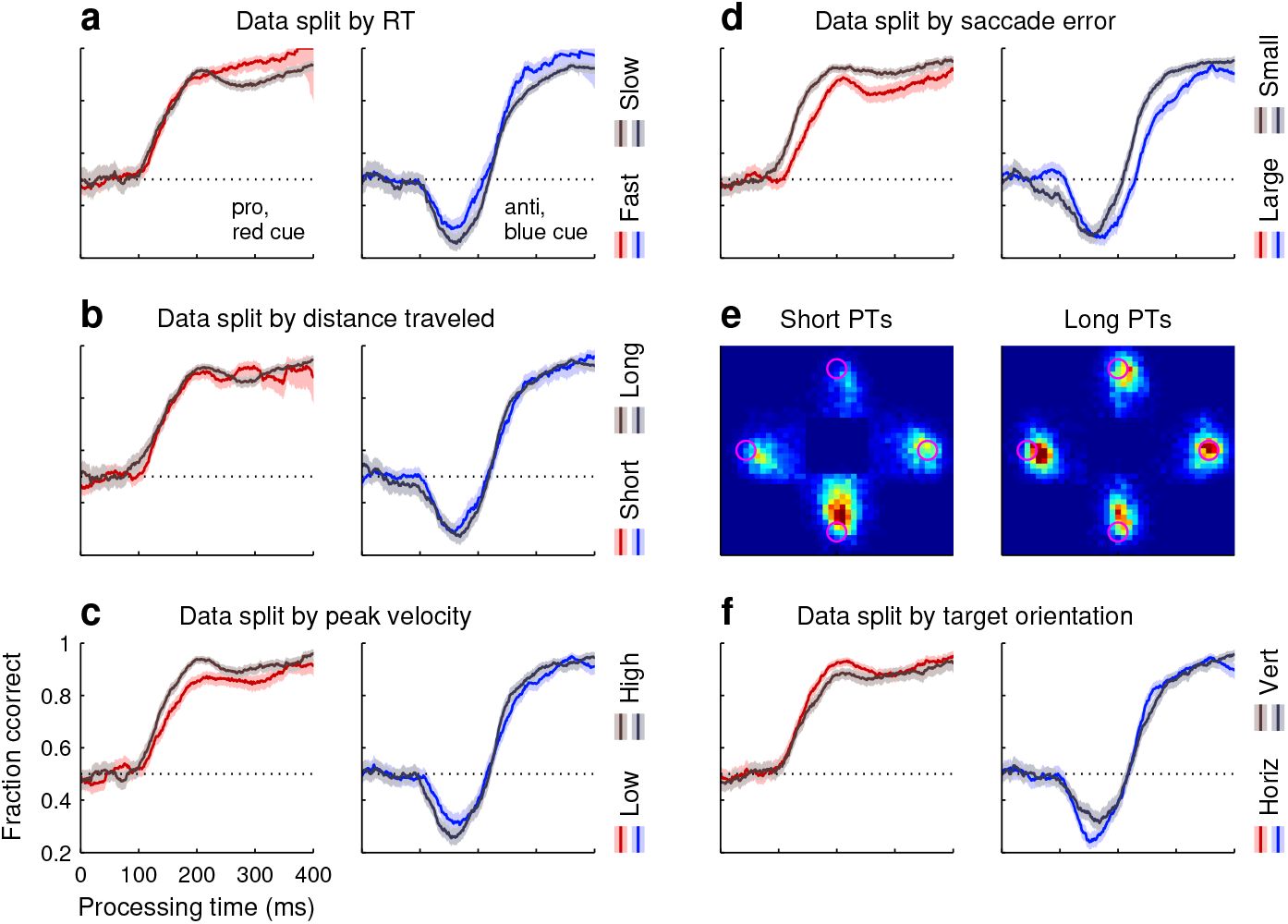
Performance in pro and anti choices correlates weakly with urgency and saccade-specific parameters. **a**, For each participant, choices from the base version of SpotChase were split according to RT into fast (below the median RT) and slow (above the median RT). The data were then combined across participants. Tachometric curves are shown for the fast-RT (saturated colors) and slow-RT (dark colors) data subsets, each subdivided into pro (red traces, left) and anti (blue traces, right) choices. Shaded ribbons denote 95% CIs. **b**, Tachometric curves for the data split according to the total angular distance traveled by the eyes during each choice (from onset of targets to 100 ms after saccade onset). **c**, Tachometric curves for the data split according to the peak velocity of the choice saccades. **d**, Tachometric curves for the data split according to the landing error of the choice saccades. **e**, Spatial density maps for saccade landing points during guesses (left, PT *<* 100 ms) and fully informed choices (right, PT *>* 250 ms). Color indicates saccade counts in each 0.5^°^×0.5^°^spatial bin, from 0 (dark blue) to maximum (red). Magenta circles indicate target locations at 8^°^ of eccentricity. **f**, Tachometric curves for the data split according to the orientation of the choice targets.

There were some evident differences in behavior between SpotChase and comparable trial-based tasks used in past studies (Salinas et al., 2019; Goldstein et al., 2022, 2024; Oor et al., 2023). For instance, in SpotChase, the PT distributions were wider in both directions, i.e., both guesses (PT ≲ 90 ms) and highly informed choices (PT ≳ 300 ms) were more frequent. To assess the consequences of such higher variance, we examined the degree to which perceptual performance correlated with three saccade-specific features: the total angular distance traversed by the eyes in each choice (from onset of targets to 100 ms after saccade onset), the peak velocity of the choice saccade, and the saccade error, which is the distance between the saccade landing point and the chosen target. First we considered inclusion/exclusion of data based on these quantities. For each of them, no appreciable effect on the tachometric curves was observed when we excluded from analysis the choices at the tails of the distribution (10% most extreme values on either the left side, right side, or both; not shown). Next, for each saccade feature, we divided the data into two subsets of equal size using a median split, as described above for the RT, and computed pro and anti tachometric curves for each subset. Results based on the total distance traveled by the eyes (mean of short distances: 14°, long distances: 22°) revealed no appreciable difference (Fig. 2b). Results based on peak saccade velocity (mean of low velocities: 313°/s, high velocities: 454°/s) revealed only a slight difference during pro choices (Fig. 2c). And results based on saccade error (mean of small errors: 1.28°, large errors: 3.36°) revealed a clear difference whereby more accurate movements were associated with perceptual judgements that were elaborated slightly more rapidly and reached a higher performance ceiling (Fig. 2d, note relative shift of ∼25 ms between rising trajectories). Such association between saccade accuracy and perceptual accuracy is consistent with prior findings (Seideman et al., 2018; Wollenberg et al., 2020).

Finally, we note a marked asymmetry in the spatial distribution of choice saccades produced during SpotChase: when targets were arranged vertically, participants generally made many more downward than upward saccades, especially when they were guessing (Fig. 2e, left panel; probability of looking at bottom target = 0.766). This asymmetry decreased with increasing PT and largely disappeared for fully informed vertical choices (PT *>* 250 ms; Fig. 2e, right panel; probability of looking at bottom target = 0.510). These results are consistent with previously reported visual asymmetries generally indicating higher perceptual sensitivity in the lower visual field compared to the upper (Himmelberg et al., 2023). The current data demonstrate a motor preference aligned with such perceptual gradient. By contrast, horizontal choices were nearly balanced between left and right targets (probability of looking at left target was 0.457 during guesses and 0.502 during fully informed choices). Importantly, though, in spite of the obvious difference between vertical and horizontal choices, their corresponding tachometric curves were nearly indistinguishable (Fig. 2f). This indicates that the relationship between processing time and success probability is robust to inherent perceptuo-motor biases.

In summary, our analyses yielded some quantitative differences across data subsets, but they were slight, indicating that oculomotor parameters relate only weakly to perceptual performance, as expected (Seideman et al., 2018). The pro and anti tachometric curves always bore the same qualitative relation to PT and to each other regardless of how the data were partitioned. Thus, overall, the results so far show that, despite the relaxed task control and highly dynamic conditions, performance in SpotChase is robust and eminently consistent with that observed during trial-based urgent tasks: in both cases perceptual accuracy is fundamentally determined by the PT, the salience of the cue onset leads to reliable oculomotor capture early on, and this exogenous (involuntary) draw can be overcome slightly later by an endogenous (voluntary) signal that is congruent with the task goals.

### Early non-random responses are salience-driven

Prior studies with urgent tasks indicate that the pronounced dip below chance observed during anti choices is due to involuntary, exogenous capture that is driven by stimulus salience and, for the most part, acts independently of endogenous goals or task rules (Goldstein et al., 2022; Oor et al., 2023). To determine whether this was indeed the case under dynamic conditions, 9 of our 19 participants performed the salient-noncue (**SNC**) variant of SpotChase (Methods).

The SNC variant followed the same layout as the base, but the luminances of the stimuli were different (Fig. 3a). Recall that, in the base version, the cue was more salient than the opposing target (noncue) because of its sudden color change. In the SNC version, the initial yellow targets were of low luminance. Then, after a variable time period, one target (cue) switched to a low-luminance red or blue while the other (noncue) turned to a high-luminance yellow. The idea was for the noncue, which was still behaviorally uninformative, to become more salient than the cue, which was still behaviorally informative.

**Figure 3.**
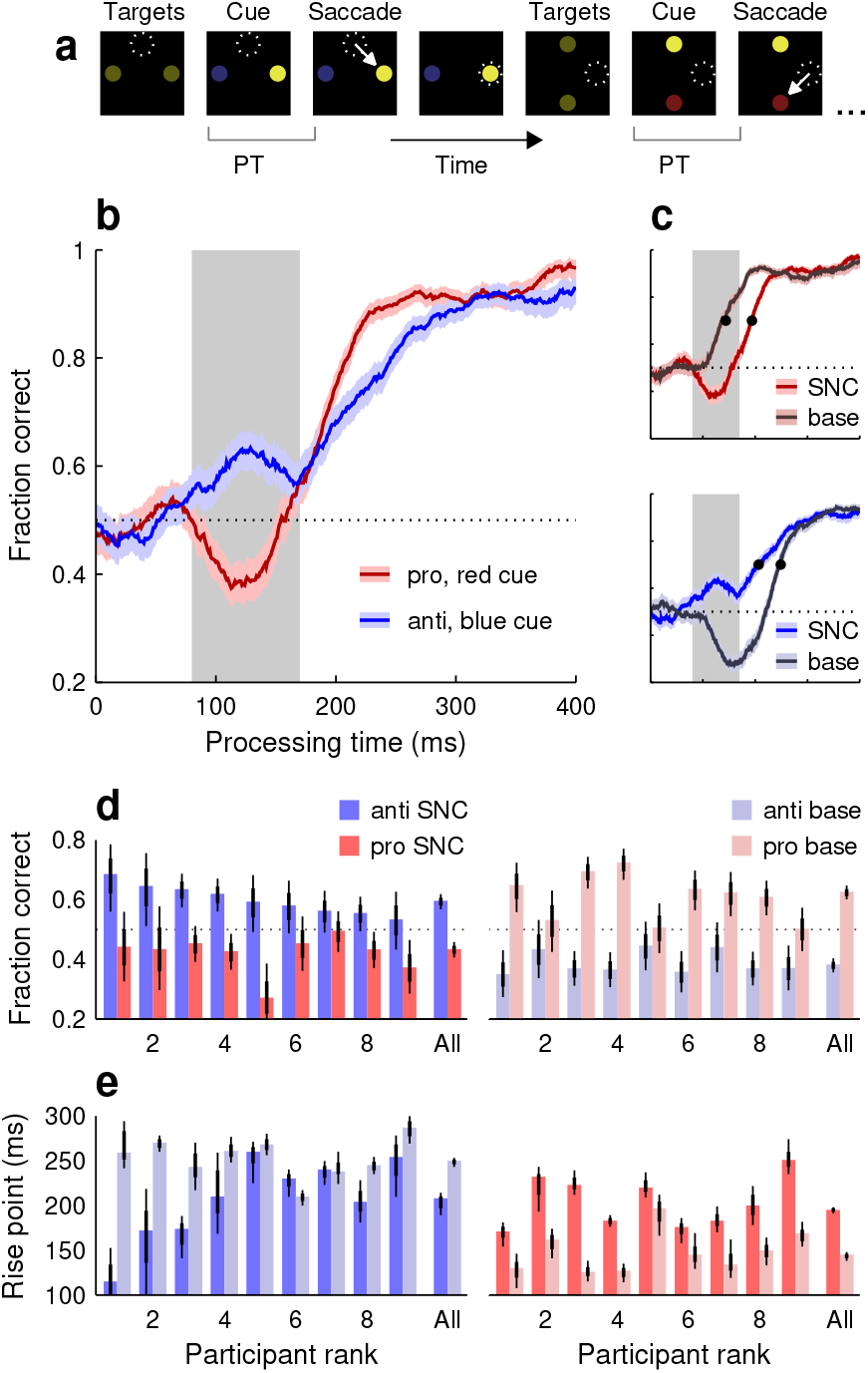
Performance in the salient-noncue (SNC) variant of SpotChase. **a**, Sequence of events in the task. The initial yellow targets are of low luminance. Following a variable time, one target (cue) turns red or blue and remains of low luminance, whereas the other (noncue) stays yellow but shifts to high luminance. The task is otherwise the same as in the base version. **b**, Tachometric curves for pro (red trace) and anti choices (blue trace) pooled across 9 participants. Shaded ribbons indicate 95% CIs across choices. Shaded rectangle marks the exogenous capture window used for analysis (80 ≤PT ≤170 ms). Horizontal dotted line indicates chance level. **c**, Comparison between tachometric curves in the SNC (bright colors) and base (dark colors) variants. Same conventions as in **b**. Upper and lower graphs show pro (red traces) and anti curves (blue traces), respectively. Data for all curves are from the same participants. Black dots mark curve rise points. **d**, Fraction of correct responses within the exogenous capture window for individual participants. Bars indicate results for pro (red) and anti (blue) choices in the SNC (saturated colors) and base conditions (light colors). Thin and thick lines mark 95% and 68% CIs, respectively. Participants are sorted by their effect magnitude in the anti SNC case. **e**, Curve rise points across participants. Same conventions and participant order as in **d**.

As before, separate tachometric curves were generated for pro and anti choices (Fig 3b). Now, the early deviations from chance (PT ≳ 80 ms) were reversed relative to the trends seen in the base condition. During anti choices (Fig. 3b, blue trace), performance showed an initial, transient uptick above chance consistent with exogenous oculomotor capture toward the salient noncue, i.e., toward the correct response. Conversely, during pro choices (Fig. 3b, red trace), performance showed precisely the opposite, a brief dip. This downward deflection was again consistent with exogenous capture toward the noncue, but in this case it corresponded to the incorrect response.

To quantify this exogenous pull, for each participant we computed the fraction correct based on all the pro or anti choices found within a restricted exogenous capture window (80 ≤PT ≤170 ms; Fig. 3b, c, gray shades; Methods). Results were highly uniform across our sample (Fig. 3d): whereas in the base task performance in this window was consistently higher for pro than for anti choices (all light red bars higher than light blue bars), in the SNC variant the opposite was true, performance was consistently higher for the anti choices (all bright blue bars higher than bright red bars). Reversing the direction of salience reliably reversed the early deflections in performance.

Additionally, the SNC results showed that the timing of the later, endogenously driven rise toward asymptotic accuracy was also impacted by the exogenous signal. Whether the correct choice was toward the cue (Fig. 3c, top graph) or toward the noncue (Fig. 3c, bottom graph), an exogenous pull away from the correct choice tended to delay the endogenously driven rise. For pro choices, the rise point for the pooled data increased by 50 ms as the salience signal shifted from the cue to the noncue (rise point in base version: 144 ms in [138, 148] ms, 95% CI; rise point in SNC version: 194 ms in [191, 198] ms, 95% CI; Fig. 3c, top, black dots). And for anti choices, the rise point increased by 42 ms as the salience signal shifted from the noncue to the cue (rise point in SNC version: 207 ms in [189, 214] ms, 95% CI; rise point in base version: 249 ms in [243, 253] ms, 95% CI; Fig. 3c, bottom, black dots). These delays were also highly consistent across individual participants (Fig. 3e; note all bright red bars above light red bars, and light blue bars above bright blue bars in 7 of 9 cases). When the salience signal was misaligned with the goal, it took longer to elaborate a correct informed choice.

These characteristic effects correspond to the simultaneous occurrence of two phenomena that have been typically considered separately. The early deviations from chance performance represent involuntary saccades toward a salient stimulus, and are akin to traditional oculomotor capture (Theeuwes et al., 1998, 1999). In turn, the delays in reaching a performance criterion represent a time cost on voluntary saccades toward a goal, and are akin to traditional attentional capture (Ruz and Lupiáñez, 2002; Theeuwes, 2010; Luck et al., 2021). These overt and covert forms of exogenous capture, which may be understood as distinct manifestations of the same underlying mechanism (Salinas et al., 2019; Salinas and Stanford, 2021; Goldstein et al., 2022; Oor et al., 2023), occur together and quite robustly under dynamic, less constrained viewing conditions.

### Advanced knowledge of cue location engages endogenous attention

One of the most characteristic aspects of visuospatial attention is that it can be deployed voluntarily and covertly, independently of gaze (Eckstein et al., 2004; Ignashchenkova et al., 2004; Thompson et al., 2005; Busse et al., 2008; Zhou and Thompson, 2009; Herrington and Assad, 2010; Carrasco, 2011). This has been amply demonstrated in tasks that require sustained fixation. To investigate the impact of endogenous, covert attention on saccadic choices under less constrained conditions, we designed a variant of our dynamic task in which participants could potentially benefit from deploying attention to specific locations in advance of their eye movements.

In the spatial bias (**SB**) variant of SpotChase (Fig. 4a; Methods), participants were made aware, before the start of each run, that the red or blue cue would always appear either at the top and left positions (SB1 runs), or at the bottom and right positions (SB2 runs). Participants were not instructed to perform any differently; they were simply informed of the spatial regularity. This way, they had the opportunity to covertly attend to the informative cue locations early on — but note that this did not give away what the correct choice was, because the cue color (red or blue) remained unpredictable.

**Figure 4.**
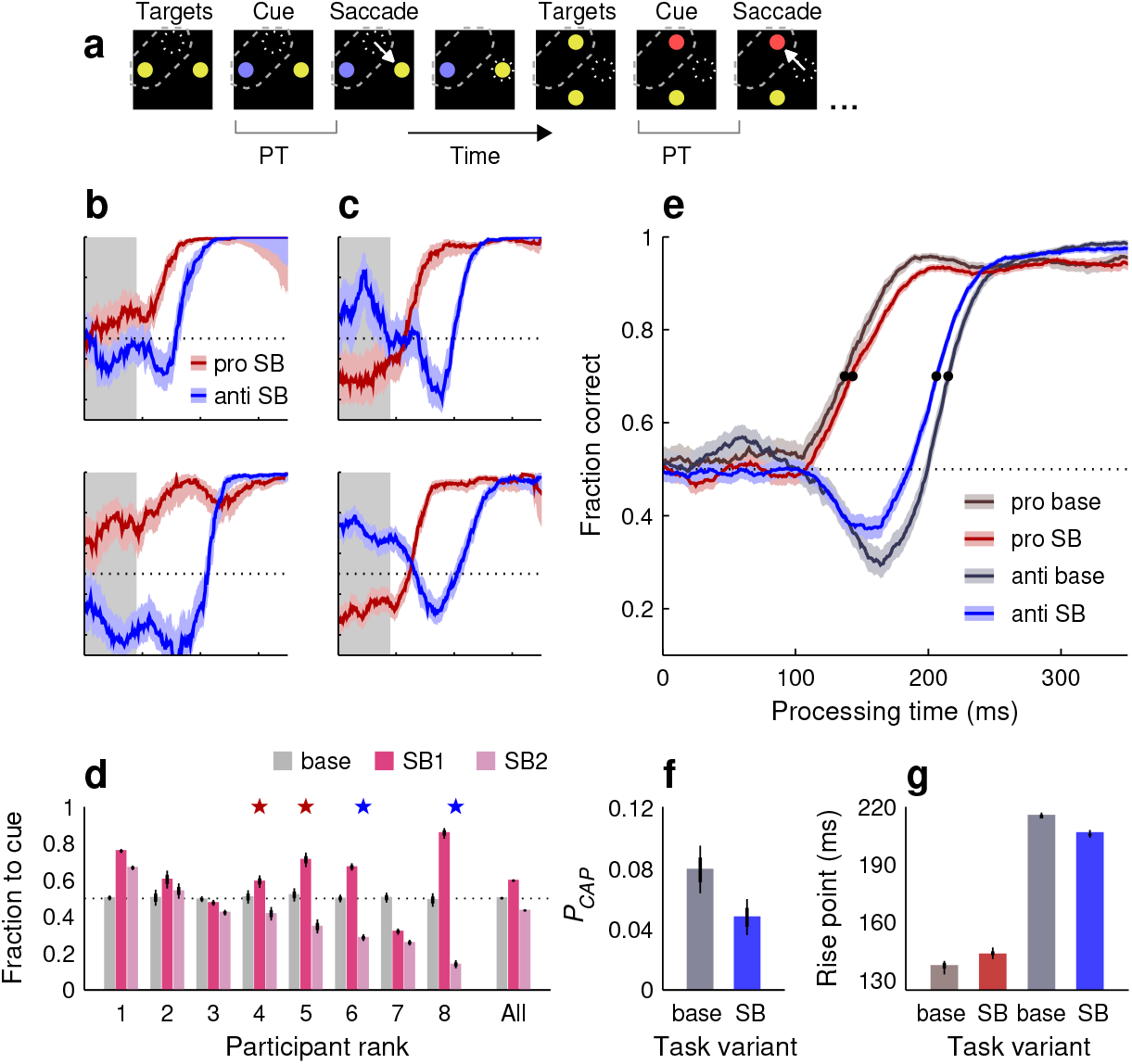
Performance in the spatial-bias (SB) variant of SpotChase. **a**, Sequence of events in the task. Participants knew that the informative red or blue cues would always appear at fixed locations, top or left in this example (SB1; dashed outlines, not shown on screen). Otherwise, the task proceeded as in the base version. **b**, Tachometric curves from two participants who demonstrated a bias toward the known cue location in the SB1 case. See panel **e** for axis labels. Shaded ribbons indicate 95% CIs across choices. Shaded rectangles mark the response-bias window (PT ≤90 ms). **c**, As in **b**, but for two participants whose responses were biased away from the known cue location in the SB2 case. **d**, Fraction of uninformed choices (guesses) toward the cue location for each participant; PT window used is indicated in **b, c**. Bars depict results when the cue appeared unpredictably (base, gray), when it appeared predictably at the top or left locations (SB1, dark pink), and when it appeared predictably at the bottom or right locations (SB2, light pink). Participants are sorted by their effect magnitude in the SB2 case. Stars mark the example participants in **b** (red pair) and **c** (blue pair). **e**, Comparison between pooled tachometric curves in the SB (saturated colors; SB1 and SB2 data combined) and base (dark colors) task variants from the same participants. Black dots mark curve rise points. **f**, Probability of capture for the anti curves shown in **e. g**, Rise points for the four curves shown in **e**. In **d, f, g**, thin and thick lines mark 95% and 68% CIs, respectively.

There is an important methodological clarification to make about this manipulation. Initial SB data were collected after the participating individuals (8 subjects) had completed the base and SNC variants of the task. In this case, some of the differences between the base and SB results could be conceivably explained as a consequence of increased practice rather than differences in attentional allocation, so we decided to repeat the SB experiment. For the subsequent replication, 8 new participants were recruited who performed interleaved runs of the base, SB1, and SB2 variants of SpotChase (group 3 participants; Methods), thus eliminating differences in practice between these conditions. In what follows, we discuss the results obtained with the replicate dataset. We note, however, that the results with the initial (sequential) and the later (interleaved) datasets were highly consistent qualitatively.

We were interested in two questions. The first one was whether the participants’ eye movements would demonstrate some form of spatial coupling to the attended locations. This issue is important for understanding the neural basis of visuospatial attention (Maunsell, 2015; Moore and Zirnsak, 2017), and springs from prior studies indicating that the voluntary deployment of attention is generally accompanied by an incipient motor plan to the attended point (Kustov and Robinson, 1996; Belopolsky and Theeuwes, 2009; Hanning et al., 2022; Goldstein et al., 2024). It is possible that such coupling becomes more obvious when strict fixation requirements are lifted (Belopolsky and Theeuwes, 2012).

Indeed, the participants’ eye movements did change quite noticeably when the cue locations were known: their uninformed choices became strongly biased. Some participants tended to make more guesses to the cue (Fig. 4b), whereas others tended to make more in the opposite direction, to the noncue (Fig. 4c). To quantify this bias, we calculated the mean fraction of choices toward the cue in the guessing range of processing times (PT ≤90 ms). By this measure, guesses in the SB variants were typically biased for any given participant (Fig. 4d; fraction to cue was significantly different from 0.5 with *p <* 10^−4^ in 14 of 16 cases), much more so than in the base condition (Fig. 4d; mean absolute deviation from 0.5 in base condition: 0.006; in SB conditions: 0.171; *p* = 0.0002, permutation test). Some participants clearly showed opposite trends in the SB1 and SB2 variants (Fig. 4d, participants 4, 5, 6, 8). On average, the tendencies for looking to the cue or to the noncue were approximately equal across the SB1 and SB2 datasets (Fig. 4d, All), so the bias in the pooled data was small (Fig. 4e, bright red and blue traces).

We interpret these results as follows. Attending to a particular location automatically creates an incipient motor plan to shift gaze toward that location. This would explain a propensity to look at the cue. On the other hand, although attending to the cue location may be perceptually advantageous, moving prematurely toward it is not because it simply results in a fast guess. An effort to prevent involuntary capture would explain a propensity to look away from the cue. Although the overall pattern of results may seem odd, it is largely consistent with a previous study in which similar pro-anti choices were made within a trial-based urgent task (Goldstein et al., 2024). In any case, knowledge of the cue location certainly had an impact on the viewing behavior of the participants. The varied directions and magnitudes of their guessing biases indicate that, although attentional deployment and saccade planning are strongly associated under dynamic conditions, their coupling is flexible and idiosyncratic.

The second question of interest in the SB experiment was whether knowledge of the cue location could bolster perceptual performance, which is the defining signature of spatial attention. Although the spatial regularity provided no information about which choice was correct, attending to the cue location early on could potentially save some processing time by expediting the identification of the cue color, for instance. On the other hand, because all the colored stimuli were of high luminance and easily discriminable from each other, we also contemplated the possibility that performance in this variant of the task, as assessed by the tachometric curves, might be the same as in the base.

The results indicate that yes, on average, covertly attending to the cue locations did bolster perceptual processing specifically during anti choices (Fig. 4e, blue curves). In comparing the pooled data from the same participants during the base and SB versions, we found that the pro tachometric curve rose 6 ms later when the cue location was known (Fig. 4e, g, red traces and bars). In contrast, the anti tachometric curve rose 9 ms earlier (Fig. 4g, blue bars; rise point in base: 215 ms in [214, 217], 95% CI; rise point in SB: 206 ms in [204, 208]). More notably, the fraction of captured saccades also decreased substantially in the SB case (Fig. 4f; *P*_*CAP*_ in base: 0.079 in [0.064, 0.095], 95% CI; *P*_*CAP*_ in SB: 0.048 in [0.036, 0.060]). These results are consistent with weaker exogenous capture and faster recovery when attention could be covertly deployed to the cue.

In summary, given advanced knowledge of relevant spatial locations, the participants exhibited widely varying motor biases indicative of strong yet flexible coupling between attentional deployment and oculomotor planning. Moreover, we also observed a perceptual benefit in the form of faster processing and weaker capture during anti choices. The results show that the vol-untary allocation of attention to relevant (cue) locations may provide a perceptual advantage even when stimuli change rapidly and are highly discriminable.

### Perceptual performance is strongly swayed by stimulus statistics

Expectations and statistical regularities in the environment can exert profound influence on perception and attention (Sterzer et al., 2008; Seriès and Seitz, 2013; de Lange et al., 2018; Rungratsameetaweemana and Serences, 2019; Klink et a., 2023). To assess the effects of stimulus (red/blue cue) or task (pro/anti) statistics on visuomotor performance, ten participants completed a new variant of SpotChase, one with variable pro-anti ratio (**PAR**; Methods). In the PAR version, red and blue cues were still presented randomly but one color was sampled more frequently than the other. The ratio of pro-to-anti (or red:blue) choices was 3:1, 6:1, 1:3, or 1:6, and each ratio yielded a full dataset from each of the ten participants. We generally expected that performance would be somewhat better for the overrepresented choice type than for the underrepresented; however, given an earlier study where only minimal differences were observed between blocked and interleaved pro/anti trials (Goldstein et al., 2022), strong effects were not anticipated.

Actually, the impact of stimulus/task statistics was surprisingly large, both for pro (Fig. 5a, b) and anti choices (Fig. 5c, d). In general, relative to the base condition (1:1 ratio), performance improved substantially when a given choice type occurred more often and deteriorated substantially when it occurred rarely, but unique trends were observed in each case. During pro choices, the early rise in performance was unaffected by the task statistics, as quantified by the rise point of the pooled tachometric curves (Fig. 5a, black dots; Fig. 5f, red bars). However, at later PTs (170 ≤PT ≤350 ms), when choices are informed by the task rules, the fraction correct reached 0.96 as red cues became more common and fell to around 0.67 as they became more rare (Fig. 5e). In contrast, during anti choices, the pooled tachometric curves shifted markedly according to the frequency of the blue cues (Fig. 5c, black dots; Fig. 5f, blue bars), with the rise point going as low as 196 ms (in [195, 198] ms, 95% CI) for the 1:6 ratio and as high as 309 ms (in [301, 319] ms, 95% CI) for the 6:1 ratio. The task statistics determined not only the timing of the rise in anti performance, but also the proportion of captured saccades, which varied dramatically as well (Fig. 5g).

**Figure 5.**
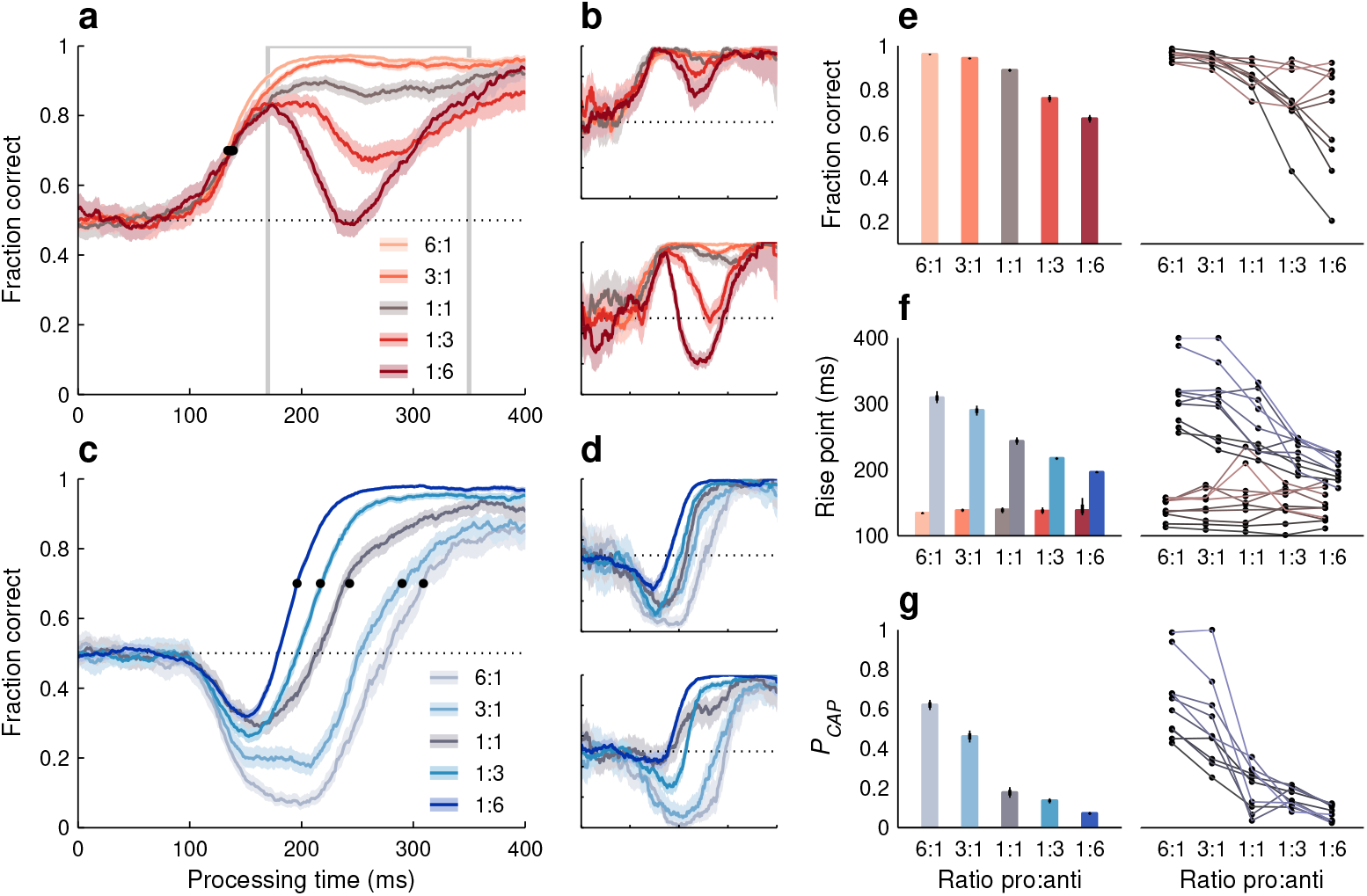
Performance in the variable pro-anti ratio (PAR) variant of SpotChase. **a**, Tachometric curves for pro (red-cue) choices pooled across participants. Each curve corresponds to a different pro:anti ratio, as indicated. Shaded ribbons indicate 95% CIs across trials. Outlined gray rectangle marks the endogenous-response window (170 ≤PT ≤350 ms). Horizontal dotted line indicates chance level. Black dots mark curve rise points. All curves are from the same 10 participants. **b**, As in **a**, but for two individual participants. **c**, Tachometric curves for anti (blue-cue) choices pooled across participants. Same conventions as in **a. d**, As in **c**, but for two individual participants. **e**, Fraction of correct pro choices in the endogenous response window (outlined in **a**) for each pro:anti ratio. **f**, Rise points for the pro (red bars) and anti (blue bars) tachometric curves for each pro:anti ratio. **g**, Probability of capture during anti choices for each pro:anti ratio. In **e**– **g**, results from pooled curves (left, bars) are shown with 95% and 68% CIs (thin and thick vertical lines). Results from individual-participant curves (right, dots) are shown with a line connecting the data from each participant. All 1:1 data are from the base version of the task.

Trends for individual participants (Fig. 5e–g, right panels) were highly consistent with those in the aggregate data (left panels). However, an interesting pattern was visible in all the quantitative measures that were sensitive to task statistics: the variability across subjects was always much smaller when the mean performance was high than when it was low. For instance, during endogenously guided pro choices (Fig. 5e), all participants achieved a fraction correct above 0.92 with the 6:1 ratio, whereas with the 1:6 ratio, the fraction correct varied widely (range: [0.2, 0.92]) around a lower mean (of 0.68). A similar distinction was observed during anti choices, both for *P*_*CAP*_ (Fig. 5g) and for the curve rise point (Fig. 5f, blue bars and lines). Thus, individual differences based on PT were strongly amplified when task conditions became more demanding.

### Sensitivity to task statistics is inconsistent with top-down control

In the PAR variant of SpotChase, participants performed better on a given choice type when it was more common. We considered two mechanisms potentially responsible for such effect. The first is top-down modulation. In this scenario, performance varies between pro-anti ratios because participants recognize a dominant choice type and strategically prioritize it at the expense of the alternative type. That is, if they notice that pro choices are more frequent, they willfully deploy attentional or cognitive resources so that pro choices become more successful, and similarly for anti choices. To test whether participants could voluntarily trade accuracy in one task against the other in such a way, the same 10 subjects who completed the PAR variant went on to perform the extreme-value (**EV**) variant of SpotChase. In this variant, everything was identical to the base version, including the 1:1 red-blue ratio, except that the point values assigned to each colored choice were different (Fig. 6a; Methods). In one case (EVB runs) blue-cue errors were much more costly than red-cue errors, whereas in another case (EVR runs) the reverse was true.

**Figure 6.**
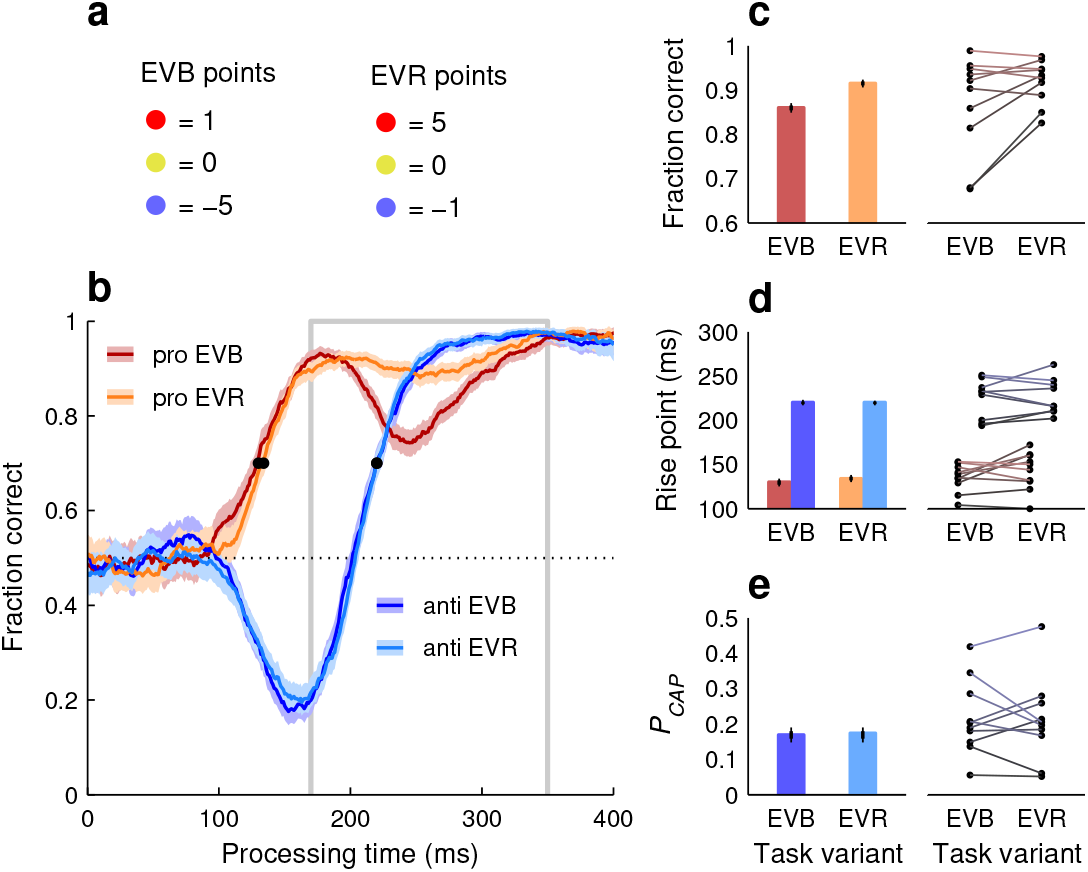
Performance in the extreme-value (EV) version of SpotChase. **a**, Point values assigned to each cue color in the EVB (anti choices prioritized) and EVR (pro choices prioritized) task variants. **b**, Pooled tachometric curves for pro (red) and anti (blue) choices in the EVB (dark colors) and EVR (light colors) variants. Shaded ribbons indicate 95% CIs across choices. Outlined gray rectangle marks the endogenous-response window (170 ≤PT ≤350 ms). Black dots mark curve rise points. **c**, Fraction of correct pro choices in the endogenous response window (outlined in **b**). **d**, Rise points for pro (red) and anti (blue) tachometric curves. **e**, Probability of capture during anti choices. In **c**–**e**, results from pooled curves (left, bars) are shown with 95% and 68% CIs (thin and thick vertical lines). Results from individual-participant curves (right, dots) are shown with a line connecting the data from each participant. All data are from the 10 participants who performed the PAR and EV variants.

Participants responded to the different incentives as intended; specifically, their overall success rate for anti choices was higher during EVB runs (68.7% correct in [67.8, 70.0], 95% CI, pooled data) than during EVR runs (65.1% in [64.2, 66.1], 95% CI), whereas their overall success rate for pro choices did not change (74.6% in both cases). This, however, largely resulted from adjustments in urgency (pace); participants slowed down when prioritizing anti choices (EVB runs: RT = 354 ±1.2 ms, mean ±SE, pooled data; EVR runs: 330 ±1.1 ms). In terms of perceptual processing capacity, as quantified by the tachometric curve, the two priority settings yielded only minimal differences (Fig. 6). The pooled anti curves were essentially identical between EVB and EVR conditions (Fig. 6b, d, e, blue traces and bars), and comparisons based on individual participant data yielded no significant differences in *P*_*CAP*_ or rise point (*p >* 0.6 in both cases, permutation test; Fig. 6d, e, dots joined by bluish lines). For the pooled pro curves, the early rise in performance was also the same across conditions (Fig. 6b, d, red/orange traces and bars). The only notable effect was a drop in the fraction correct in the later endogenous range (170 ≤PT ≤350 ms) when anti choices were prioritized (EVB case). This decrease in performance was evident for the pooled data and clear (no overlap between 95% CIs) for 5 of the 10 participants (Fig. 6c).

In summary, participants were severely limited in their ability to voluntarily sway their perceptual performance in favor of one specific choice type or the other. Most notably, the prioritization of anti choices (EVB case) did not improve the processing of the corresponding blue cues in any way relative to the opposite criterion (EVR case). This means that willful intent cannot explain the strong variations in performance observed in the PAR variant (Fig. 5). Therefore, such variations must have resulted fundamentally from involuntary mechanisms.

### Sensitivity to task statistics is consistent with selection history

Given the preceding results, we considered the role of selection history effects. Specifically, we examined how recent experience (within a few past choices) related to the variations in performance due to longer-term experience (within many runs) that resulted from the different ratio conditions. To this end, the data from each ratio in the PAR experiment were split by choice type (pro or anti) and sorted based on the history of cue colors that preceded each choice (Methods). We focused on histories denoted as 2R, 1R, 1B, and 2B, in which a choice of interest was immediately preceded by 2 red cues, 1 red cue, 1 blue cue, or 2 blue cues, respectively. For example, with this nomenclature, a 2R pro choice is one in which the current cue is red and the two preceding cues were also red; similarly, a 1R anti choice is one in which the current cue is blue and the preceding cue was red; and so on. After sorting the data in this way, a history-conditioned tachometric curve was constructed for each combination of ratio and history considered.

We found that the effects of short- and long-term experience — set by history and ratio conditions — were consistent with each other, and together had a profound impact on perception (Fig. 7). In general, performance for a given cue color improved both when that color was more frequent overall and when it was repeated from the previous choices to the current one; and conversely, performance for a given cue color deteriorated when such color was less frequent and when it represented a switch relative to the color in the previous choices.

**Figure 7.**
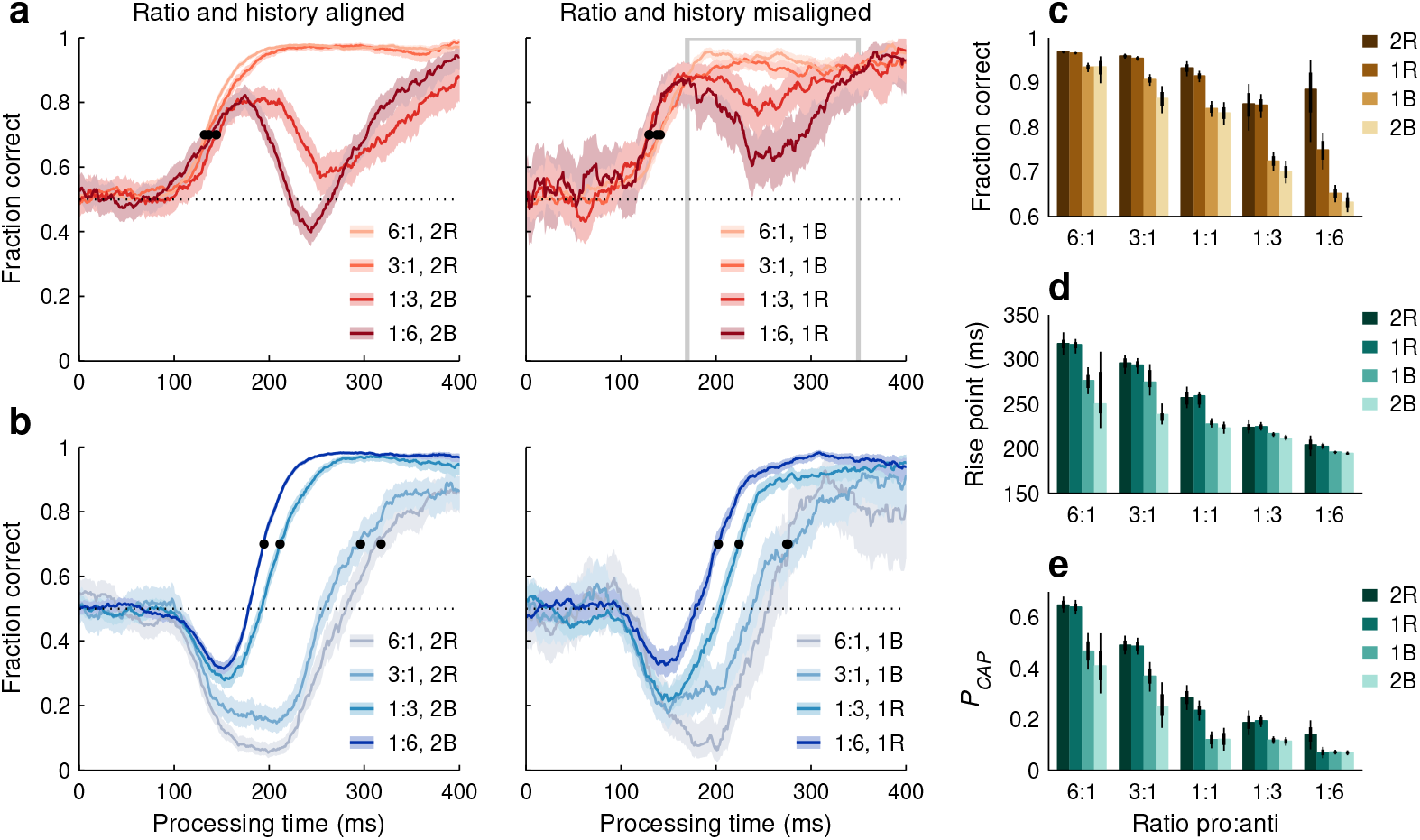
SpotChase performance depends on both short- and long-term experience. For each pro:anti ratio value, data were pooled over participants, and pro and anti choices were conditioned on the prior history of cue colors (sequences 2R, 1R, 1B, 2B). **a**, Tachometric curves for pro choices. The left panel shows results conditioned on aligned ratio and history combinations, in which both factors yield a high incidence of prior red cues (6:1, 3:1 ratios, R histories) or blue cues (1:6, 1:3 ratios, B histories). The right panel shows results conditioned on misaligned combinations, in which one factor yields higher and the other lower incidence of a given cue color. Other conventions as in Fig. 5a. **b**, As in **a**, but for anti choices. **c**–**e**, Behavioral metrics as functions of pro:anti ratio and history sequence. Bars indicate the fraction of correct pro choices in the endogenous response window (**c**), and the rise point (**d**) and probability of capture (**e**) of the anti tachometric curve. Of the 20 values shown in each panel, 8 correspond to the tachometric curves shown in **a** or **b**. Thin and thick lines indicate 95% and 68% CIs.

For pro choices (Fig. 7a, c), large variations in success rate occurred during the endogenous response window (170 ≤PT ≤350 ms). These were largest when the ratio and history conditions reinforced each other (Fig. 7a, left; aligned conditions), i.e., when they both augmented or dimin-ished the incidence of prior red cues. In this case, the fraction correct for a pro choice went from 0.97 when red cues were most common and the two prior choices presented two red cues (6:1, 2R case), to 0.63 when red cues were least common and the two prior choices presented two blue cues (1:6, 2B case). In contrast, when the ratio and history conditions conflicted with each other (Fig. 7a, right; misaligned conditions), their effects on performance tended to cancel out. In this case, the variations in fraction correct were considerably smaller, even though the data spanned the same set of ratios. The fraction correct for a pro choice went from 0.93 when red cues were most common but the prior choice presented a blue cue (6:1, 1B case), to 0.75 when red cues were least common but the prior choice presented a red cue (1:6, 1R case).

During anti choices (Fig. 7, b, d, e), performance was similarly influenced by both short- and long-term experience. Relative shifts in the pooled tachometric curves were largest again when the ratio and history conditions reinforced each other (black points in Fig. 7b, left; aligned conditions), either both augmenting the incidence of prior blue cues (yielding best performance) or both diminishing it (yielding worst performance). Here, the anti rise point spanned 123 ms between the most and least favorable combinations (rise point in 1:6, 2B case: 195 ms in [193, 197] ms, 95% CI; in 6:1, 2R case: 318 ms in [305, 330] ms). Notably, when the ratio and history conditions were in conflict (Fig. 7b, right; misaligned conditions), the pooled curves became closer to each other, now spanning only 73 ms between the two extremes (in 1:6, 1R case: 203 ms in [199, 207] ms; in 6:1, 1B case: 276 ms in [261, 291] ms). The same ratio and history combinations yielded similarly conspicuous differences in the proportion of captured saccades (Fig. 7e).

To quantify and more easily visualize the relative contributions of short- and long-term experience to perceptual performance, the three behavioral metrics derived from the pooled tachometric curves were arranged as joint functions of ratio and history (Fig. 7c–e). For this analysis, the data from the PAR and base (1:1 ratio) variants were combined. Qualitatively, the three metrics used to characterize performance demonstrated the same three patterns. (1) Relative to a recent switch, one recent color repetition consistently led to better performance. This can be seen by comparing the 1R and 1B results: for pro choices, 1R always produced a higher fraction correct than 1B (Fig. 7c, no overlap in 95% CIs for all 5 cases), whereas for anti choices, 1B typically produced lower *P*_*CAP*_ and rise point values than 1R (Fig. 7d, e, no overlap in 95% CIs for 8 of 10 cases). (2) The results were generally similar for one versus two repetitions of a given color (Fig. 7c–e, overlap in 95% CIs in 14 of 15 cases for both 2R versus 1R bars and 2B versus 1B bars). Taking into account two trials into the past was only slightly more predictive than taking into account only one (Fig. 7c–e, differences between 2R and 2B values tended to be larger than differences between 1R and 1B values; *p >* 0.06, permutation tests). (3) In all three metrics, there was substantial variation both across histories and across ratios (*p <* 0.002 for history, *p <* 3 ×10^−5^ for ratio, two-way ANOVAs). However, the different ratios consistently accounted for a larger percentage of the variance in the results (Fig. 7; in panel c: 66% and 25% due to ratio and history; in panel d: 84% and 12%; in panel e: 77% and 16%).

We interpret these patterns as follows. The consistent influence of immediate history (R vs B contrasts) reveals the action of some adaptation mechanisms that are fast and local. Although such mechanisms may appear to saturate rapidly when considering only a few choices into the past (2R vs 1R, and 2B vs 1B contrasts), this does not imply the absence of slower mechanisms that respond to more global variations in stimulus statistics (ratio contrasts), and whose effects may go largely undetected unless the analyses can go farther into the past. Thus, the full-blown impact of past events on a current choice becomes evident only when the task statistics are manipulated to permit reliable comparisons across both long- and short-term histories.

The history of prior cue colors produced large variations in performance in SpotChase. Now, given that reward (or success) is a potent driver of behavior and modulator of attention (Milstein and Dorris, 2007; Chelazzi et al., 2013; Failing and Theeuwes, 2018; Oor et al., 2025), we also asked: was the history of prior outcomes (correct or incorrect) also relevant to such variations? The answer is essentially “no.” The PAR data were again split by choice type (pro or anti), but this time they were sorted according to outcome history. We considered all the choices of a given type that were preceded by one correct choice (1C), two correct choices (2C), etc., or by one error (1E), two errors (2E), etc. The resulting tachometric curves conditioned on outcome history largely overlapped each other, and the three metrics used for characterizing them demonstrated minimal variations across sequences. Analyses combining both outcome and cue-color histories did not reveal a substantial effect of outcome either. Thus, the trial-by-trial history effects were fundamentally feature-based.

### Experience-driven effects characterize individual variability

After manipulating the stimulus/task statistics, we noted that, when the task became more difficult, performance varied widely between participants (Figs. 5e–g). This was true for both pro and anti choices, and we initially thought that the two effects would be consistent; that is, that each participant would demonstrate a particular sensitivity to history regardless of choice type. Interestingly, however, the dependencies of pro and anti choices on recent experience turned out to be unrelated.

For each participant who performed the PAR variant of SpotChase, two quantities were calculated to describe their sensitivity to past cue-color exposure, a Δ fraction correct (during pro choices) and a Δ*P*_*CAP*_ (during anti choices). Each Δ was equal to the (positive) difference between values in the most favorable and least favorable ratio-history combinations considered earlier. That is, the data from each participant were analyzed as in Fig. 7c, d, and for each of the two behavioral metrics considered, the difference between the two extreme conditions (6:1, 2R and 1:6, 2B) was taken. This yielded a Δ fraction correct and a Δ*P*_*CAP*_ for each participant. The pairing of these two quantities produced data points that were widely spread (Fig. 8a). For some partic-ipants, both values were indeed small (e.g., E1) or large (e.g., E4), indicating similarly weak or similarly strong sensitivity to past cue colors during pro and anti choices alike. But overall there was no correlation; for other participants (e.g., E2, E3), the modulation was large in one direction but small in the other.

**Figure 8.**
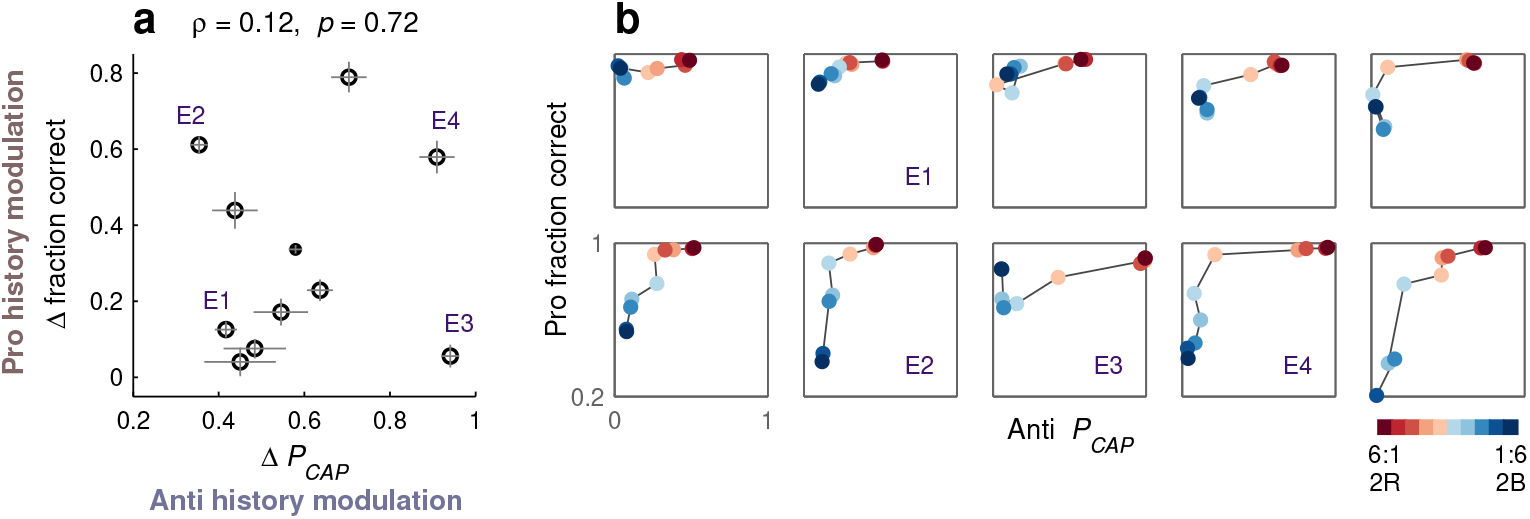
History-driven effects characterize individual variability. **a**, Sensitivity to cue-color history in pro choices (Δ fraction correct) versus in anti choices (Δ*P*_*CAP*_). Each circle corresponds to one participant. Error bars indicate ±1 SE. For reference, the small point shows results for the pooled data. Pearson correlation coefficient and significance (from permutation test) are indicated. **b**, History-driven tradeoffs between pro and anti performance. Each square contains data from one participant. Each colored point shows the fraction correct in pro choices (y axis) versus the probability of capture (*P*_*CAP*_) in anti choices (x axis) for a given ratio-history combination. Colorbar indicates depicted combinations favoring pro (red spectrum: 6:1, 2R; 6:1, 1R; 3:1, 2R; 3:1, 1R; 1:1, 1R) or anti (blue spectrum: 1:1, 1B; 1:3, 1B; 1:3, 2B; 1:6, 1B; 1:6, 2B) choices. Labels correspond to the four example participants marked in **a**.

The range of individual sensitivities becomes even more evident when the fraction correct from pro choices and *P*_*CAP*_ from anti choices are plotted against each other to visualize how they change together across different ratio-history combinations (Fig. 8b). For this analysis, we considered 10 of the 20 combinations used in Fig. 7c–e (all the aligned combinations plus two additional; see Fig. 8, caption), for which performance varied systematically and over a wide range. In the resulting representation, the dark blue point is always close to the left y-axis, indicating best anti performance (*P*_*CAP*_ near zero), whereas the dark red point is always close to the top x-axis, indicating best pro performance (fraction correct near 1). But, notably, the trajectory from one to the other varies between participants denoting various possible tradeoffs between pro and anti performance. Some participants exhibit only a moderate drop in anti performance when red cues are highly likely (E1). Others show a much larger variation in pro than in anti performance (E2), or vice versa (E3). And in some cases, performance plummets whenever the encountered cue color, either red or blue, is experienced infrequently (E4).

These results indicate not only that sensitivity to recent history is a major source of individual variability in perceptual capacity, but also that there are multiple, distinct adaptation mechanisms that make independent contributions to such variability.

## Discussion

SpotChase produces viewing conditions that are more dynamic and less constrained than in traditional trial-based tasks, yet it permits rigorous quantification of perceptual capacity. This revealed clearly identifiable contributions to visuomotor performance from stimulus-driven, goal-driven, and history-driven attention mechanisms within a continuously flowing behavior.

Even highly established phenomena can change unpredictably in less stringent paradigms that mimic naturalistic settings (Camerer and Mobbs, 2017; Cohen et al., 2020; Snow and Culham, 2021). Here, the most unexpected result was that history-driven involuntary mechanisms exerted a surprisingly powerful influence on perceptual performance by modulating the relative probability of success of pro and anti choices We also found that the interplay between stimulus-driven and goal-driven mechanisms was qualitatively similar to that observed in urgent, trial-based tasks — but the corresponding data were also interesting. For instance, they showed that there are benefits to the covert allocation of attention even when visual scenes change rapidly, consist of highly discriminable stimuli, and yield idiosyncratic saccade planning.

To appreciate the results, it is worth stressing the fundamental distinction between the RT, which is anchored to the start of motor planning (here, onset of targets), and the PT, which is anchored to the start of perceptual processing (here, cue onset). Whereas the RT is a direct reflection of urgency, i.e., how rapidly motor plans advance, the PT is not (Fig. 2a). A key consequence of this is that the behavioral curve that describes choice accuracy as a function of PT, the tachometric curve, can be read as a representation of how the probability of success changes over time *during a single choice* (Salinas et al., 2010; Stanford et al., 2010; Costello et al., 2013; Seideman et al., 2018; Salinas et al., 2019; Oor et al., 2023). That is, although the tachometric curve is constructed by sorting different trials according to PT, modeling and neurophysiological results indicate that its shape is a faithful record of the sequence of neural events that unfold within each choice.

According to this interpretation, at the time at which accuracy first departs systematically from chance, near the 100 ms mark, the target selection circuitry receives an exogenous input signaling the cue onset. This input has all the hallmarks of an involuntary bias: it is fast, it tracks the most salient stimulus regardless of task rules (Fig. 3b), and its effects are uniform across individuals (Fig. 3d, e). It can be thought of as a transient, stimulus-driven pulse of neural activity that automatically biases ongoing motor plans toward salient, recently detected targets (Bisley et al., 2004; Ipata et al., 2006; Scerra et al., 2019; Goldstein et al., 2022; Klink et al., 2023; Zhu et al., 2024). Later on (PT ≳ 170 ms), motor plans are steered toward the correct choice by an endogenous signal that also depends on the cue, but after it has been interpreted according to task rules. This signal can change substantially across task conditions and individual participants (Figs. 3b, 5, 7).

Thus, between 100 and 350 ms after cue onset, the dynamics of attention control and target selection are remarkably lawful and rich. Indeed, distinctive neurophysiological markers of attentional processing occur in this range (Luck, 2012). Resolving these dynamics psychophysically is key for relating behavior to neural activity. Specifically, whereas the time point at which a neural circuit discriminates two spatial locations can be determined with high accuracy (∼tens of ms), the equivalent time point for the behavioral judgement generally has a much larger margin of error (Bisley and Goldberg, 2003; Busse et al., 2008; Cohen et al., 2009; Herrington and Assad, 2010). When the measurement is based on PT, however, the two can be directly compared at high resolution (Seideman et al., 2022; Zhu et al., 2024).

In experiments lacking time pressure, psychometric functions of time have been generated by varying the presentation duration of stimuli, often subsequently masked, and measuring report performance at each duration (e.g., McAvinue et al., 2012; Habekost et al., 2013). The steepness of the resulting curve, which represents information accrual per unit time, is then considered a measure of perceptual processing capacity. This interpretation applies to the tachometric curve as well, but two qualifications are important. First, based on limited stimulus duration, one can never be sure how long the sensory signal circulated in the decision-making circuitry and was effectively available for processing. In contrast, the PT derived under urgent conditions eschews this ambiguity, so estimates of processing speed based on the tachometric curve have higher temporal precision (for a detailed example, see Seideman et al., 2022, Supplementary Note). And second, the steepness of a curve will depend on exactly what is being processed and how (e.g., Poth and Schneider, 2024). Specifically, in an urgent, trial-based version of a simple prosaccade task (Goldstein et al., 2022), we observed an extraordinarily steep rise that took only a few milliseconds. Several factors likely contributed to such extreme processing speed: the cue appeared alone, was highly discriminable, and only had to be detected, because the response type (pro) was known before the cue onset. In contrast, the pro curves in SpotChase rose more gradually, but the conditions were more complex: the cue was shown alongside another stimulus (noncue) that altered its exogenous draw, and its color had to be resolved to determine the correct motor response.

In general, pro choices are processed rapidly because the endogenous and exogenous signals are aligned, in which case they are typically difficult to dissociate (Aagten-Murphy and Bays, 2017; Goldstein et al., 2022). Here, the PAR and EV variants of SpotChase provided particularly clean evidence of their separate contributions by revealing that the later rise in accuracy (PT .:2 170 ms) may vary dramatically with history (Fig. 7a, b, e) and, to a lesser degree, with top-down priority (Fig. 6b–d). The distinctions between exogenous and endogenous signals were stark. Normally, however, whenever an intended saccade target is salient, the informed, goal-driven signal seamlessly prolongs the effect of the earlier stimulus-driven pulse, so target selection appears to be guided by a single continous process.

When the exogenous pulse of activity is not aligned with the rule-based signal (anti choices), it produces either overtly captured, erroneous saccades or correct saccades with processing delays (Figs. 1, 3) — effects analogous to traditional oculomotor and attentional capture (Theeuwes et al., 1998, 1999; Ruz and Lupiáñez, 2002; Luck et al., 2021; Theeuwes, 2010), as mentioned earlier. Such effects are not exclusively tied to physical salience. For instance, Poth (2021) reported qualitatively similar findings using urgent versions of classic cognitive tasks in which physical salience plays no role (a spatial Stroop task and a flanker task). We suspect that capture phenomena are the signatures of a more general and ubiquitous type of conflict, that between low-level (rapid, easily available) and high-level (slower, computationally expensive) signals (Kahneman, 2011; Aagten-Murphy and Bays, 2017; Poth, 2021; Krause and Poth, 2025). These signatures are fully revealed as functions of PT.

Pointing to broader generality beyond the realm of visual perception, the results are also consistent with experiments on cognitive control and conflict adaptation (Egner, 2017; Poth, 2021), which show that recent history is typically a strong modulator of cognitive performance. According to this literature, whenever top-down control is necessary to overcome a habitual response that would lead to an incorrect choice (e.g., the Stroop task), performance is highly sensitive to both the short- and long-term incidence of conflict trials. That is, performance on a conflict trial is typically better when the previous trial was also a conflict trial (Egner, 2017), and when the longer-term frequency of conflict trials is high (Bugg, 2017). If the current data are indeed representative of this trend, it would imply that the importance of history effects has, in fact, been underestimated. The results show that history-driven modulation is strongest at critical processing times during which the probability of success changes rapidly (here, 100 ≲ PT ≲ 350 ms). In contrast, non-urgent tasks can only detect the residual differences seen afterward, once this probability has approached its ceiling — but these late differences are much weaker. Perhaps the same is true for task switching more generally (Wylie and Allport, 2000; Monsell, 2003). We suggest that, if performance were more often pushed to its temporal limits, the importance of selection history to cognitive processing would become more obvious for many types of task.

The dynamic, uninterrupted viewing conditions of SpotChase likely created an ideal environment for history-driven effects to manifest. These were complex, in that they involved short and long time scales, varied widely across individual participants, and seemed to consist of separate, independent components associated with pro and anti choices. Relative to most prior studies documenting experience-driven biases on perceptual performance, the observed variations were huge, especially given that the stimuli were highly discriminable (for a detailed discussion of PT-dependent history effects compared to the wider literature, see Oor et al., 2025). The data, however, were in line with a recent study (Adam et al., 2024) demonstrating that a salient distracter presented for the very first time produces much stronger attentional capture than in typical experiments in which conditions are repeated. Expectation and/or recent experience matters, indeed.

## Materials and Methods

### Subjects and setup

Experimental participants were 27 healthy human volunteers (17 females, 11 males; median age: 28 years; age range: 21–57 years) recruited from the Wake Forest Baptist Medical Center and broader Winston-Salem communities. All participants had normal or corrected-to-normal vision and provided written informed consent prior to the experiment. All experimental procedures were approved by the Institutional Review Board of Wake Forest University School of Medicine in accordance with protocol IRB 00035241.

Similar to a previous study (Goldstein et al., 2022), all experimental sessions were conducted in a dimly lit room. Participants were seated on an adjustable chair with their forehead and chin supported. Stimuli were presented using a VIEWPixx LED monitor (VPixx Technologies Inc., Saint Bruno, Quebec, Canada; 1920 x 1200 screen resolution, 120 Hz, refresh rate, 12-bit color) placed 57 cm in front of the participant. Eye position was monitored monocularly and recorded using an EyeLink 1000 infrared camera and tracker (SR Research, Ottawa, Canada; 1000 Hz sampling rate). All stimulus presentation and data collection routines were implemented using Matlab (Mathworks, Natick, MA) and the Psychtoolbox 3.0 package (Brainard, 1997; Kleiner, 2007) running on a standard Windows-based desktop computer. Psychtoolbox incorporates the Eyelink Toolbox (Cornelissen et al., 2002) for control of the eye tracker.

### Experimental design and statistical analyses

#### BASE TASK

Every variant of SpotChase followed the same general sequence of events as the base version (Fig. 1a), which proceeded as follows. Data were collected in “runs,” each one lasting 90 seconds. A run began with the participant fixating on a red target at the top-center of the screen. Following fixation, two yellow targets (RGB vector [0.62 0.61 0], 1.5° in diameter) appeared at 8 ° to the left and right of the screen center. This was followed by a variable interval referred to as the stimulus onset asynchrony, or SOA (50–300 ms in increments of 50 ms). After the SOA elapsed, one yellow target, referred to as the cue, switched to either red (RGB vector [1 0.25 0.25]) or blue (RGB vector [0.25 0.55 1]). All stimuli were of high luminance (32 cd m^−2^, where the black background had a luminance of 0.04 cd m^−2^). Thereafter, nothing changed until the participant made a saccade into one of the two invisible spatial boxes surrounding the cue and the noncue. Such boxes were rectangular, each centered on the corresponding target stimulus, and measured 7×10°, with the long side aligned with the choice axis (horizontal, in this case). Following a choice, i.e., a saccade into one of the target boxes, the display remained the same for an additional 150 ms. Then, simultaneously, the horizontal targets disappeared and two vertical yellow targets appeared at 8° up and down from the screen center. Thus, the choice cycle started anew, unfolding with the same sequence of events: targets on, cue, and choice saccade. Horizontal and vertical choices alternated thereafter throughout the duration of the run. Cue colors and locations were selected randomly.

The instructions given to participants were simple: to look at one of the two target stimuli present on the screen at any point in time, in order to maximize their score at the end of the run. Participants could make saccadic choices before or after the cue was revealed. The run score depended on the point values assigned to the three stimulus colors: 3 points were awarded for looking at a red target, 1 for looking at a yellow target, and 0 for looking at a blue target. To incentivize participants to make choices rapidly, even if that meant guessing on occasion, in a small fraction of the choices (10%) the cue was never shown (infinite SOA). In this case, participants had to select one of the two yellow targets at random. This got the participants accustomed to guessing and discouraged them from waiting too long for the cue. The actual score for each run was computed using the total accrued points divided by the total reaction time (RT) taken by all the choices, where the RT of an individual choice was measured from the onset of the yellow targets to the onset of the choice saccade. The score depended on both the participant’s accuracy for discriminating colors and on their urgency, because given the fixed duration of each run (90 s), the less time spent pondering each choice, the more choices could be made. It corresponded to a reward rate, and was given in points per hour. Participants also received a “bonus,” which was added to their score. The bonus was equal to the total number of correct minus the total number of incorrect choices, also divided by the total RT. It was relatively small compared to the main score (∼18%), and corresponded to the chance-adjusted correct rate, in hits per hour.

#### TASK VARIANTS

To acclimate them to the task environment, participants performed a few short (30 s) practice runs, first with red cues only, then with blue cues only, and then with both interleaved. Participants required 3–5 such practice runs before beginning data collection with the base version of SpotChase. All variants of SpotChase followed the same conventions as the base (Fig. 1a), but with alterations to the properties of the displayed stmuli. In the salient-noncue variant (SNC; Fig. 3a), the initial yellow targets were of low luminance (RGB vector [0.11 0.11 0]). Then, after the SOA, the noncue switched to a high-luminance yellow and was paired with either a low-luminance red (RGB vector [0.16 0 0]) or a low-luminance blue cue (RGB vector [0 0.03 0.23]). The (low) luminance was the same for the three dim colors (5 cd m^™2^), and the (high) luminance for the bright yellow was the same as for the base variant (32 cd m^™2^). In this case, the yellow, uninformative noncue was meant to be more physically salient than the red or blue cues, which still informed what the better choice was.

In the two spatial bias (SB) variants of SpotChase (Fig. 4a), SB1 and SB2, all stimulus colors and task conditions were the same as in the base version except that the cue locations were predictable. In the SB1 case, the cue always appeared on the top (in vertical choices) or on the left (in horizontal choices); whereas in the SB2 case, the cue always appeared on the bottom (in vertical choices) or on the right (in horizontal choices). In this version of the task, for each choice, the participants knew where the informative cue would appear, but still could not predict the location of the higher-value stimulus.

The pro-anti ratio (PAR) variant of SpotChase was the same as the base version except that, instead of the red and blue cues appearing with equal probabilities (1:1 red-to-blue ratio), one of them was more frequent. Participants who performed this version of the task experienced either predominantly red cues (3:1 or 6:1 ratio) or predominantly blue cues (1:3 or 1:6 ratio), with the probability ratio fixed for any given run. For this experiment, participants completed full datasets in a fixed sequence of ratios: 3:1, 6:1, 1:3, and 1:6. In this case, relative to cue salience, one of the two stimulus-response rules (look at the salient cue; look at the less salient noncue) was much more common than the other.

Finally, in the extreme-value (EV) variants of SpotChase, the stimuli were again the same as in the base condition but their assigned point values were different. In the EVB case, red, yellow, and blue stimuli were worth 1, 0, and −5 points, respectively, so the priority for participants was to avoid looking at any blue target. In the EVR case, red, yellow, and blue stimuli were worth 5, 0, and −1 points, respectively, so the priority for participants was to look at any red target. Participants were informed of the changes in point values and made aware that the run score would be maximized by prioritizing performance either on blue-cue choices (during EVB runs) or red-cue choices (during EVR runs). Participants first completed the EVB runs and then the EVR runs.

The 27 participants belonged to three groups. All members of group 1 (7 females, 2 males) performed the base version of SpotChase first (13–17 runs) and then the SNC variant; subsequently, 8 members of group 1 (7 females, 1 male) also peformed the SB1 and SB2 variants, in that order. All members of group 2 (6 females, 4 males) performed the base version of SpotChase first, then the PAR variant, and then the EV variant. All members of group 3 (4 females, 4 males) performed the base and SB variants of SpotChase by intereleaving runs in the following order: base, SB1, SB2, and repeating the cycle 18–32 times. The first 19 participants were randomly assigned to groups 1 and 2; the remaining 8 participants, assigned to group 3, were recruited specifically to replicate the SB experiment more rigorously, i.e., to disambiguate effects that could be due to the order in which the task variants were performed. Data from group 3 are presented only in Fig. 4 and related text; all other results are from groups 1 and 2.

#### DATA PROCESSING

As mentioned above, a participant made a choice when they brought their gaze for the first time into one of the two applicable target boxes. Offline, the choice saccade was the same one detected at runtime, except that its onset was calculated using a velocity criterion (40°/s). Thus, for each choice offered, the RT was computed as the interval between the onset of the yellow targets and the onset of the corresponding response saccade into the target box. A minimum saccade amplitude was not explicitly imposed. Choices were excluded from analysis if they occurred before the onset of the yellow targets (which would produce RT *<* 0); if the RT was too long (*>* 800 ms); if a blink occurred within 100 ms of the response saccade onset; or if entry into a valid response box was registered but the velocity threshold was not met. Apart from these cases, which constituted approximately 14% of all detected choice saccades, no other choice saccades were excluded from analysis. Results were robust relative to changes in data inclusion/exclusion criteria (Fig. 2).

From each participant and each of the two main task conditions (red cue, blue cue), we collected a median of 1,122 choices (range: 584–1,431) suitable for analysis in the base version of the task. We aimed for this amount of data based on prior results indicating that this number was appropriate for resolving individual performance changes with a temporal resolution on the order of 10 ms or so (Shankar et al., 2011; Salinas et al., 2019; Goldstein et al, 2022, 2024). The numbers were similar for the other task variants, except for the ratio or PAR variant. In that case, we aimed to collect approximately the same amount of data (∼1,100 choices) from the less frequent condition, so many more trials were collected from the complementary, high-frequency one. For example, for a pro-anti ratio of 3:1, the 1,100 anti choices were accompanied by approximately 3,300 pro choices.

Except for the very start of each run, no explicit fixation requirements were imposed. Participants sometimes made a saccade to the target opposite to the chosen one immediately after making an error, and sometimes made a saccade to the center of the screen before making an unequivocal choice. These movements, and others that did not bring the eyes into the boxes surrounding the two current valid targets, were not considered response saccades and were ignored. Critically, their existence was not detrimental to the measured relationship between choice accuracy and time: the PT was measured from the moment the color cue was revealed until the eyes reported an unambiguous choice, and the PT consistently dictated the probability of success regardless of whether intermediate (non-choice) saccades were more or less likely (Fig. 2b).

The PT or cue viewing time associated with each choice was equal to the time elapsed between the onset of the cue and the onset of the response saccade (which entered one of the target boxes). In practice, it was calculated as PT = RT −SOA, where all three quantities are specific to each choice. For analysis purposes, a choice was scored as ‘correct’ when the participant looked at the target with the higher assigned value given the alternatives, even if the corresponding saccade was initiated when both targets were still yellow (i.e., it was a guess).

#### DATA ANALYSIS

Data analyses were performed in the Matlab programming environment (The MathWorks, Natick, MA). As in the case of trial-based urgent tasks (Salinas et al., 2019; Poth, 2021; Goldstein et al., 2022, 2024; Oor et al., 2023, 2025), tachometric curves were produced by indexing trials by PT and sorting them into PT bins that shifted every 1 ms. The bin width was 81 ms for calculating the probability of capture (explained below) and 41 ms for all other analyses. The fraction of correct responses was then calculated for all the trials inside each bin. The resulting curve describes how choice accuracy depends on PT, or cue viewing time.

Whenever possible, analyses were carried out both for individual participants and for their pooled data. Pooling means that all the choices from a given group of participants were combined into one large dataset, as if produced by a single, aggregate participant.

For some analyses, performance was quantified as the mean fraction of correct choices over a wide PT window. For the SB variant, choice biases were measured via the fraction correct for guesses (PT *<* 90 ms). For the SNC variant, we considered the fraction correct in an intermediate window where exogenous capture was most likely (80 *<* PT *<* 170 ms). For the PAR variant, we considered the mean fraction correct at a long-PT window for which choices were most likely informed by the cue (170 *<* PT *<* 350 ms). For any given fraction correct, a 95% or 68% confidence interval (CI) was calculated using binomial statistics; specifically, the Agresti-Coull method (Agresti and Coull, 1998).

To quantify the magnitude of exogenous capture, we measured the fraction correct within a fixed PT window, as described in the previous paragraph, and we also calculated the probability of capture, or *P*_*CAP*_, which does not require a predetermined PT window. It was defined as

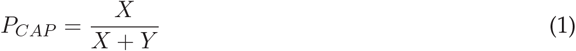

where *X* represents the area between the tachometric curve and the chance line when the curve is below said line, and *Y* is the area between the curve and the chance line when the curve is above. This quantity corresponds to the proportion of erroneous choices that cannot be ascribed to chance: of those saccades that were not guesses, *P*_*CAP*_ is the fraction that were directed toward the worse alternative. Thus, *P*_*CAP*_ equals 0 when performance is at or above chance for the full PT range considered, and equals 1 when performance dips below chance at some point and never goes above. Shown results are based on *P*_*CAP*_ as computed directly from the empirical tachometric curves in the range 70–400 ms. An alternative calculation of *P*_*CAP*_, also defined via Equation 1 but derived from analytical fits to the tachometric curves, produced qualitatively similar results. CIs for *P*_*CAP*_ were obtained by bootstrapping (see below).

We also designed a randomization test specifically to evaluate the significance of each *P*_*CAP*_ value, i.e., the probability that a positive value was obtained just by chance when, in fact, performance was never truly below 50% correct. To to this end, we found all the PT bins where the value of the tachometric curve dipped below 50% correct and substituted the empirical ordinate values with random binomial samples (also below 50% correct) based on the same numbers of trials contained in each PT bin and assuming that the true probability correct was 50%. Then *P*_*CAP*_ was recomputed from the resulting (resampled) tachometric curve. This resampling process was repeated 10,000 times to obtain a distribution consistent with the (null) hypothesis that the performance curve dipped below chance only because of random fluctutations due to limited sampling. The significance p-value was equal to the fraction of resampled *P*_*CAP*_ values that was equal to or larger than that obtained empirically; it had a precision of 0.0001, given the number of iterations used.

As in prior studies (Shankar et al., 2011; Salinas et al., 2019; Goldstein et al., 2022, 2024), the empirical tachometric curves were fitted with continous analytical functions to extract key performance parameters. Tachometric curves that increased monotonically (e.g., for red-cue choices in the base task) were fitted with an increasing sigmoid function defined as

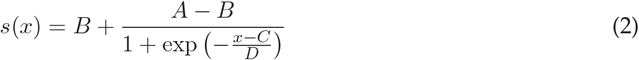

where *D* determines the slope of the curve; *B* and *A* represent chance and asymptotic accuracy levels, respectively; and *C* is the PT at which the fraction correct is halfway between *B* and *A*.

Tachometric curves that showed a substantial dip below chance (e.g., during anti choices in the base task) were fitted using a combination of two sigmoidal curves, as previously described (Salinas et al., 2019; Goldstein et al., 2022). The fitting curve *v*(*x*) was defined as

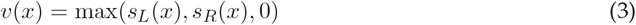

where the function max(*a, b, c*) returns the largest of *a, b*, or *c*. Here, the left and right sigmoids are given by

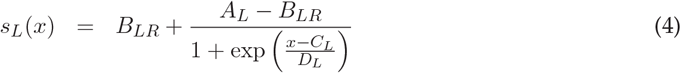

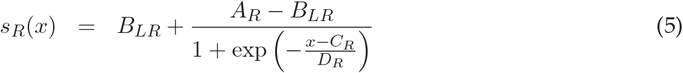

where the left, decreasing side of the tachometric curve is tracked by *s*_*L*_ and the right, increasing side is tracked by *s*_*R*_. Here, *A*_*L*_ represents chance performance (at short PTs), *B*_*LR*_ represents minimum performance (below chance), and *C*_*L*_ is the halfway point between the two. Similarly, *A*_*R*_ represents asymptotic accuracy (at long PTs), and *C*_*R*_ denotes the halfway point between the minimum (*B*_*LR*_) and asymptotic (*A*_*R*_) levels.

The main quantity derived from the analytical fits was the rise point, which was defined as the PT at which the fitting curve first exceeds a threshold value of 70% correct. By using such a fixed performance criterion, the timing of pro (red-cue) and anti (blue-cue) choices could be compared most directly (black points in Fig. 1b–d). In addition, as mentioned above, the same fits were used to compute *P*_*CAP*_ in an alternative way. For this, Equation 1 was applied to each fitting curve.

Tachometric curves were fitted to Equations 2–5 by finding parameter values (*A, B, C*, etc) that minimized the mean absolute error between the empirical data and the corresponding analytical curves. The Matlab function <monospace>fminsearch</monospace> was used for this optimization step. As done previously (Salinas et al., 2019), CIs were calculated for the fit parameters, as well as for any quantities derived from the fits (rise point, *P*_*CAP*_), by bootstrapping (Davison and Hinkley, 2006; Hesterberg, 2014). That is, the trial-wise data were resampled with replacement, the resulting (resampled) tachometric curves were re-fitted, the new parameter values and derived quantities were saved, and the process was repeated many times (1000–10,000 iterations) to generate distributions for all the parameters and derived quantities. Finally, 95% CIs were obtained by calculating the 2.5 and 97.5 percentiles derived from the bootstrapped distributions.

Although the reported rise points were typically obtained via curve fitting, as described above, we also implemented a second method for computing them. It served both as a check on the fits and as an alternative for cases in which the tachometric curves were too noisy for reliable fitting. This alternative method operates on the empirical tachometric curve and consists of two steps. First, find any PTs where the tachometric curve is near 70% correct (say, between 65% and 75%). Then, from those PTs, select the largest, continuous PT interval for which (1) all the curve values are within the specified accuracy range, and (2) the curve values bordering the left and right edges of the interval are below and above the 65% and 75% limits, respectively. The rise point is then equal to the midpoint of the resulting PT interval. As with all other quantities, CIs were generated by bootstrap. This method produced very similar rise points as that based on curve fitting. It was used primarily for the history analyses, for which data were typically parsed into small subsets.

To compare different datasets where each data point represents an experimental result from one participant, significance was determined using permutation tests for paired data (Siegel and Castellan, 1988) or equivalent randomization tests for non-paired data. To compare groups involving binary data (correct vs incorrect), confidence intervals and significance values were determined using binomial statistics (Agresti and Coull, 1988). When describing results from multiple individual participants, reported significance values for single-participant tests are uncorrected for multiple comparisons.

#### DATA SORTING FOR HISTORY ANALYSIS

History-conditioned tachometric curves (Fig. 7a, b) were generated by only considering choices that were preceded by specific cue-color sequences. For this, choices were sorted post hoc according to the number of consecutive occurrences in which the cue color on immediately preceding choices was either red (R) or blue (B). Choices preceded by a red cue (in the prior choice) were denoted as 1R, whereas choices preceded by a blue cue were denoted as 1B. Similarly, choices preceded by two consecutive red cues were denoted as 2R, whereas choices preceded by two consecutive blue cues were denoted as 2B. Thus, for example, the cue-color sequence associated with a 2R pro choice is red-red-**red**, where the bolded word indicates the cue color in the current choice, whose outcome is being evaluated. Similarly, the sequence associated with a 2R anti choice is red-red-**blue**; the sequence associated with a 1B anti choice is blue-**blue**; and so on. Applied to either pro or anti choices, the four color histories (2R, 1R, 1B, 2B) give rise to eight distinct sequences. Coded this way, 1R and 2R histories reveal the impact of color repetition on pro performance and of color switches on anti performance. And conversely, 1B and 2B histories reveal the impact of color repetition on anti performance and of color switches on pro performance.

Separate history-conditioned tachometric curves were computed based on the data from the base version of the task (pro-anti ratio of 1:1) and based on the data from each ratio in the PAR variant (6:1, 3:1, 1:3, and 1:6). This effectively combined the long-term effect (set over many runs) of a given pro-anti ratio with the the short-term effect (set over one or two choices) of a given history. Across the ratio datasets, a chosen history could thus be aligned with the dominant cue color (e.g., 2R history in 6:1 ratio) or misaligned with it (e.g., 2B history in 6:1 ratio). By this we mean that conditioning on history would be expected to either reinforce or weaken the effect of task statistics on performance, either increasing (aligned) or decreasing (aligned) any departure with respect to performance in the base version of SpotChase.

## Sources

All content was generated by humans.

## Acknowledgments

Research was supported by the National Institutes of Health (NIH) through grants R01EY025172 and R21MH120784. The project was also supported by the National Center for Advancing Translational Sciences (NCATS), NIH, through grant UM1TR004929. The content is solely the responsibility of the authors and does not necessarily represent the official views of the NIH. We thank Denise Anderson for expert technical assistance.

## Abbreviations

CI: confidence interval
EV: extreme-value
PAR: pro-anti ratio
*P*_*CAP*_: probability of capture
PT: processing time
RT: reaction time
SB: spatial bias
SNC: salient noncue
SOA: stimulus onset asynchrony

